# The predicted bZIP transcription factor ZIP-1 promotes resistance to intracellular infection in *Caenorhabditis elegans*

**DOI:** 10.1101/2021.06.17.448850

**Authors:** Vladimir Lažetić, Fengting Wu, Lianne B. Cohen, Kirthi C. Reddy, Ya-Ting Chang, Spencer S. Gang, Gira Bhabha, Emily R. Troemel

**Affiliations:** Division of Biological Sciences, University of California, San Diego, La Jolla, California, United States; Department of Cell Biology, New York University School of Medicine, New York, United States

## Abstract

Defense against intracellular infection has been extensively studied in vertebrate hosts, but less is known about invertebrate hosts. For example, almost nothing is known about the transcription factors that induce defense against intracellular intestinal infection in the model nematode *Caenorhabditis elegans.* Two types of intracellular pathogens that naturally infect the *C. elegans* intestine are the Orsay virus, which is a positive-sense RNA virus, and microsporidia, which are fungal pathogens. Surprisingly, these molecularly distinct pathogens induce a common host transcriptional response called the Intracellular Pathogen Response (IPR). Here we describe *zip-1* as an IPR regulator that functions downstream of all known IPR activating and regulatory pathways. *zip-1* encodes a putative bZIP transcription factor of previously unknown function, and we show how *zip-1* controls induction of a subset of genes upon IPR activation. ZIP-1 protein is expressed in the nuclei of intestinal cells, and is required in the intestine to upregulate IPR gene expression. Importantly, *zip-1* promotes resistance to infection by the Orsay virus and by microsporidia in intestinal cells. Altogether, our results indicate that *zip-1* represents a central hub for all triggers of the IPR, and that this transcription factor plays a protective role against intracellular pathogen infection in *C. elegans*.

## Introduction

Viruses and other obligate intracellular pathogens are responsible for a myriad of serious illnesses (1). RNA viruses, like the single-stranded, positive-sense RNA virus SARS-CoV-2 that causes COVID-19, are detected by RIG-I-like receptors (2–4). These receptors detect viral RNA replication products and trigger transcriptional upregulation of interferon genes to induce anti-viral defense (5). The nematode *Caenorhabditis elegans* provides a simple model host to understanding responses to RNA viruses, as a single-stranded, positive-sense RNA virus from Orsay, France infects *C. elegans* in the wild (6). Interestingly, natural variation in *drh-1,* a *C. elegans* gene encoding a RIG-I-like receptor, was found to underlie natural variation in resistance to the Orsay virus (7). Several studies indicate that detection of viral RNA by the *drh-1* receptor induces an anti-viral response through regulating RNA interference (RNAi) (7–9).

In addition to regulating RNAi, *drh-1* detection of viral replication products was recently shown to activate a transcriptional immune/stress response in *C. elegans* called the Intracellular Pathogen Response (IPR) (10). The IPR was defined as a common transcriptional response to the Orsay virus and a molecularly distinct natural intracellular pathogen of *C. elegans* called *Nematocida parisii* (11–13). *N. parisii* is a species of Microsporidia, which comprise a phylum of obligate intracellular fungal pathogens that infect a large range of animal hosts including humans. It is not known which host receptors detects *N. parisii* infection, but the *drh-1* RIG-I-like receptor appears to detect viral RNA replication products, and to be critical for viral induction of the IPR (10). Notably, *C. elegans* does not have clear orthologs of interferon, or the signaling factors that act downstream of RIG-I-like receptors in mammals, such as the transcription factors NF-kB and IRF3/7 (14). It is unknown how *drh-1* activates the IPR transcriptional program in *C. elegans*.

Several non-infection inputs can also trigger IPR gene expression. For example, proteotoxic stress, such as that caused by blockade of the proteasome, or by prolonged heat stress, will induce IPR genes. While intracellular infection by the Orsay virus or by *N. parisii* cause hallmarks of proteotoxic stress in *C. elegans* intestinal cells (11), genetic and kinetic analysis indicates that proteotoxic stress is activating IPR gene expression in parallel to viral infection, and there are several independent triggers of the IPR (10). Another trigger of the IPR is mutation in the enzyme Purine Nucleoside Phosphorylase-1, PNP-1, which acts in *C. elegans* intestinal epithelial cells to regulate pathogen resistance and the majority of IPR genes (12, 13, 15). Of note, mutations in human PNP cause T-cell dysfunction, but its role in epithelial cells is less well-described (15). In addition to *pnp-1*, analysis of another IPR repressor called *pals-22,* has provided insight into the regulation and function of IPR genes (12, 13). *pals-22* belongs to the *pals* (Protein containing ALS2cr12 signature) gene family, which has one ortholog each in mouse and human of unknown function, while this family has expanded to 39 members in *C. elegans* (12, 16, 17). The biochemical functions of *pals* genes are unknown, but they play important roles in intracellular infection in *C. elegans* (13). Several *pals* genes (e.g. *pals-5*) are upregulated by virus infection and the other IPR triggers mentioned above. Furthermore, two *pals* genes, *pals-22* and *pals-25*, are opposing regulators of the IPR, acting as an ON/OFF switch for IPR gene expression as well as resistance to infection (13). Not only do *pals-22/25* control immunity, but they also control thermotolerance, a phenotype that is dependent on a subset of IPR genes that encode a newly described, multi-subunit, E3 ubiquitin ligase that promotes proteostasis (13, 18).

While Orsay virus infection, *N. parisii* infection, proteotoxic stress, *pnp-1* and *pals-22* mutations all appear to act independently of each other to trigger IPR gene expression, here we show that they converge on a common downstream transcription factor. Using two RNAi screens, we find that the gene encoding a putative basic region-leucine zipper (bZIP) transcription factor called *zip-1* plays a role in activating expression of the IPR gene *pals-5* by all known IPR triggers. Furthermore, we use proteasome inhibition as a trigger to show that *zip-1* controls induction of only a subset of IPR genes. These results demonstrate that there are at least three classes of IPR genes as defined by whether their induction is dependent on *zip-1* early after proteasome inhibition, late after proteasome inhibition, or their induction after proteasome inhibition is independent of *zip-1*. We show that the ZIP-1::GFP protein is induced in intestinal and epidermal nuclei upon IPR activation, and that ZIP-1 functions in the intestine to activate *pals-5* expression. Viral induction of ZIP-1::GFP expression in intestinal nuclei depends on the DRH-1 receptor, suggesting that DRH-1 controls activation of ZIP-1. Importantly, we find that *zip-1* promotes defense against viral as well as against microsporidia infection in the intestine. Altogether, our results define *zip-1* as a central signaling hub, controlling induction of IPR gene expression in response to a wide range of triggers, including diverse intracellular pathogens, other stressors, and genetic regulators. Furthermore, this study describes ZIP-1 as the first transcription factor shown to promote an inducible defense response against intracellular intestinal infection in *C. elegans*.

## Results

### Two independent screens for regulators of the IPR identify the predicted transcription factor ZIP-1

To determine which transcription factor(s) activates IPR gene expression, we screened an RNAi library composed of 363 RNAi clones targeting 357 predicted transcription factors (less than half the predicted transcription factor repertoire in *C. elegans*) to identify RNAi clones that repress constitutive expression of the PALS-5::GFP translational reporter (*jyEx191)* in a *pals-22(jy3)* background. In parallel, we also screened this library for RNAi clones that prevent induction of the *pals-5*p::GFP transcriptional reporter (*jyIs8*) upon prolonged heat stress. In both screens, we found that *zip-1(RNAi)* led to a substantial decrease in GFP signal (Fig. 1 A and B, Table S1), suggesting that this putative bZIP-containing transcription factor plays a role in IPR regulation. We confirmed this *zip-1(RNAi)* phenotype in another *pals-22* loss-of-function allele, *jy1,* showing that here too, RNAi against *zip-1* repressed the constitutive expression of PALS-5::GFP in *pals-22* mutants (Fig. 1C). To demonstrate that this phenotype is not just restricted to *zip-1(RNAi)*, we created a full deletion allele of *zip-1* called *jy13* (Fig. S1), and observed decreased expression of the *pals-5*p::GFP reporter following prolonged heat stress in this putative *zip-1* null mutant (Fig. 1D). These results indicate that *zip-1* is important for regulating expression of two different *pals-5* GFP reporters by two different IPR triggers.

**Fig. 1.**
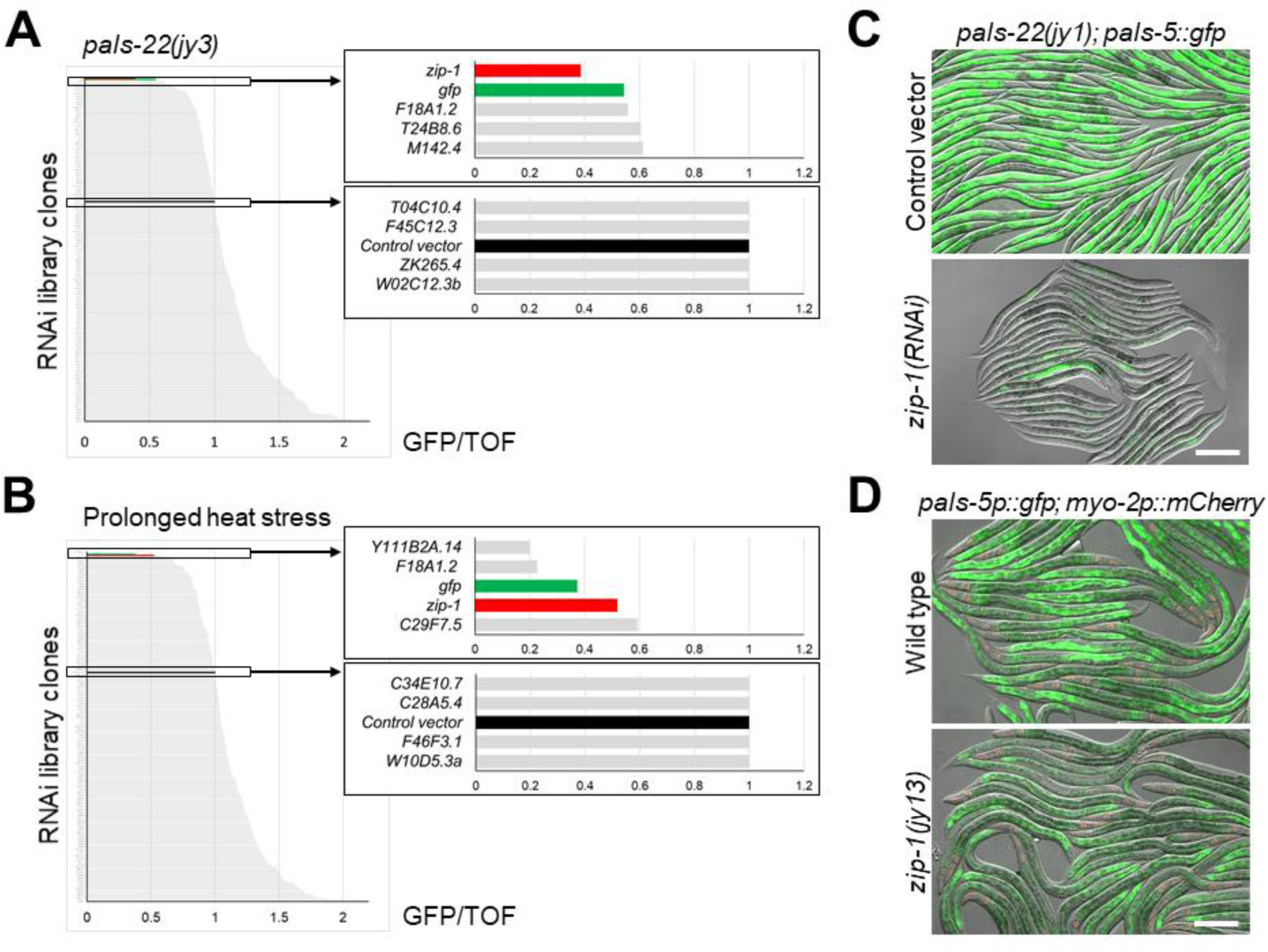
*zip-1* is required for induction of *pals-5* GFP reporters by *pals-22* RNAi and by prolonged heat stress. (A, B) Graphical overview of RNAi screen results in the *pals-22(jy3); jyEx191[pals-5::gfp]* background (A) and following chronic heat stress (B). GFP intensity was normalized to the length of worms (TOF) and it is indicated on the x-axis; different RNAi clones are listed on the y-axis. Boxes on the right represent enlarged sections of the graph containing *zip-1(RNAi)* results and relevant controls. (C) *pals-22(jy1); jyEx191[pals-5::gfp]* animals show constitutive expression of the PALS-5::GFP reporter when grown on control vector RNAi plates (upper image) but not on *zip-1* RNAi plates (lower image). (D) Expression of GFP from the *jyIs8[pals-5p::gfp, myo-2p::mCherry]* reporter is decreased in *zip-1(jy13)* animals following prolonged heat stress (lower image), in comparison to wild-type animals (upper image). (C, D) Fluorescent and DIC images were merged. Scale bars = 200 µm. *myo-2*p::mCherry is expressed in the pharynx and is a marker for the presence of the *jyIs8* transgene.

We also specifically examined whether RNAi against two other transcription factors implicated in intracellular infection were required to induce *pals-5*p::GFP expression. First, we analyzed the bZIP transcription factor *zip-10*, which is induced as part of the IPR and also promotes *N. parisii* sporulation (13, 19). Here we found no effect of *zip-10* RNAi on *pals-5*p::GFP induction upon heat stress. We also analyzed the STAT-like transcription factor *sta-2*, which is important for response to the fungal pathogen *Drechmeria coniospora*, which penetrates and grows inside epidermal cells (20, 21). Here as well, we did not find an effect of *sta-2* RNAi on induction of *pals-5*p::GFP expression (Fig. S2).

### *zip-1* is required for induction of *pals-5*p::GFP expression by all known IPR triggers

We also investigated whether *zip-1* was required for inducing *pals-5*p::GFP expression upon other IPR triggers. First, we tested whether *zip-1* was required for response to infection with the Orsay virus. Here we found that while infection of wild-type animals with the Orsay virus induced expression of the *pals-5*p::GFP reporter, infection of *zip-1(jy13)* mutants caused no GFP induction (Fig. 2A). Similarly, infection with the microsporidian species *N. parisii* caused GFP expression throughout intestine in wild-type animals, but little to no GFP expression in *zip-1(jy13)* mutants (Fig. 2B). Therefore, *zip-1* is required for induction of *pals-5*p::GFP expression after infection by these two natural intracellular pathogens of the *C. elegans* intestine. We next tested if *zip-1* was required for constitutive expression of *pals-5*p::GFP in *pnp-1* mutants (15). Here, we also saw a requirement for *zip-1*, as *zip-1(jy13); pnp-1(jy90)* double mutants had much less *pals-5*p::GFP signal compared to *pnp-1* single mutants (Fig. 2 C and D). Finally, we investigated whether *zip-1* is required for induction of *pals-5*p::GFP using the drug bortezomib (10, 11, 18, 22). Here we also saw that *zip-1* was required for induction of *pals-5*p::GFP across a timecourse of bortezomib treatment (Fig. 2 E and F). Therefore, of the six IPR triggers we tested, *zip-1* was required for induction of *pals-5p*::GFP by all of them.

**Fig. 2.**
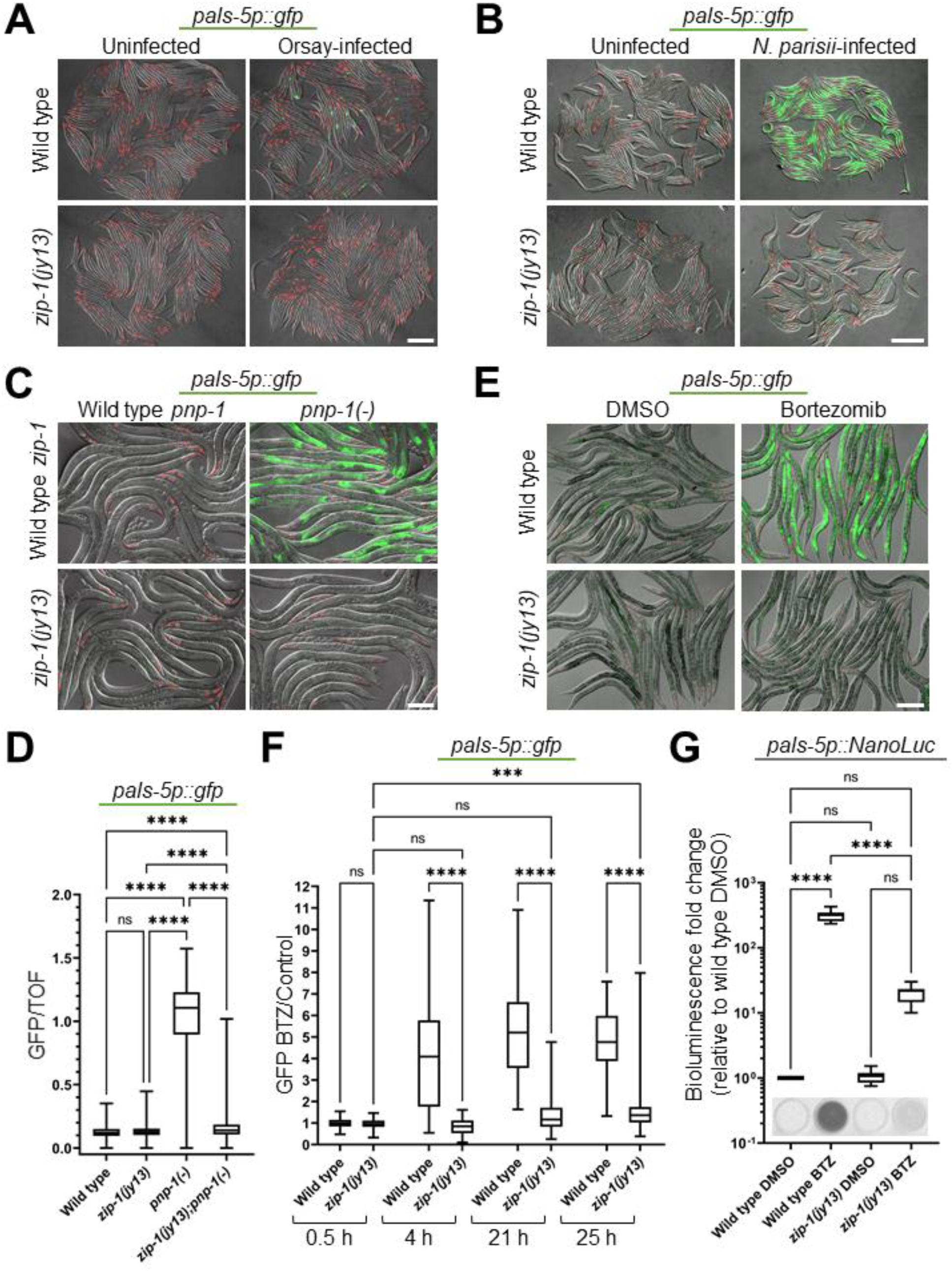
*zip-1* is required for induction of *pals-5*p::GFP expression by intracellular infections, *pnp-1* downregulation and proteasome blockade. (A, B) Intracellular infection by Orsay virus (A) and by *N. parisii* (B) leads to *pals-5*p::GFP expression in wild-type animals, but not in *zip-1(jy13)* mutants. (C) *pnp-1(jy90)* mutants show constitutive expression of the *pals-5*p::GFP reporter, which is suppressed in *zip-1(jy13); pnp-1(jy90)* double mutants. (D) Box-and-whisker plot of *pals-5*p::GFP expression normalized to length of animals (TOF). Increased GFP signal in *pnp-1(jy90)* mutants is significantly reduced in *zip-1(jy13); pnp-1(jy90)* double mutants. Three experimental replicates with 400 animals per replicate were analyzed for each strain. (E) Bortezomib treatment induces expression of *pals-5p::gfp* in a wild-type background, but not in *zip-1(jy13)* mutants. (A-C, E) Fluorescent and DIC images were merged. Scale bars = 200 µm. *myo-2*p::mCherry is expressed in the pharynx and is a marker for the presence of the *jyIs8* transgene. (F) Timecourse analysis of *pals-5*p::GFP expression in control and *zip-1(jy13)* strains following bortezomib treatment. GFP signal normalized to worm area is shown as a fluorescence intensity ratio between bortezomib-and DMSO-treated samples (y axis). Three experimental replicates with 30 animals per replicate were analyzed; average value was used for DMSO controls. Allele names and timepoints of analysis are indicated on the x axis. (G) Expression of *pals-5*p::NanoLuc reporter is significantly lower in *zip-1(jy13)* animals than in the wild-type control strain, following bortezomib treatment. Three experimental replicates consisting of three biological replicates were analyzed for each strain and treatment. Results were normalized to background luminescence and to average value of three biological replicates for wild type treated with DMSO. Normalized Relative Fluorescent Units (RLU) are shown on the y axis. Images of bioluminescent signal in representative analyzed wells are shown on the bottom of the graph. (D, F, G) In box-and-whisker plots, each box represents 50% of the data closest to the median value (line in the box). Whiskers span the values outside of the box. A Kruskal-Wallis test (D, F) or ordinary one-way ANOVA test (G) were used to calculate *p*-values; **** *p* < 0.0001; *** *p* < 0.001; ns indicates nonsignificant difference (*p* > 0.05).

Because the *jyIs8[pals-5p::gfp]* and *jyEx191[pals-5::gfp]* reporters described above are multi-copy transgene arrays that could be prone to silencing, we considered the possibility that *zip-1* repressed GFP expression in the previous experiments through its effects on transgene silencing. Therefore, we next investigated if *zip-1* inhibited expression from a single-copy transcriptional reporter, as single-copy transgenes are much less prone to silencing than multi-copy arrays. Here we used the strain with a single-copy *jySi44[pals-5p::NanoLuc]* transgene insertion, where the *pals-5* promoter drives expression of the bioluminescent protein nanoluciferase (22). Here we also found that *zip-1* was required for induction of *pals-5*p::NanoLuc bioluminescence by proteasome blockade (Fig. 2G), further indicating that *zip-1* regulates gene expression driven by the *pals-5* promoter.

Because the *zip-1* genomic locus contains a non-coding RNA *y75b8a.55* in one of its introns, and this non-coding RNA is also deleted in the *zip-1(jy13)* deletion strain, we also created a partial deletion allele of *zip-1* called *jy14* (Fig. S1). This *y75b8a.55* non-coding RNA locus is preserved in the *jy14* allele, while the region encoding the predicted bZIP domain of *zip-1* is deleted. Here with intracellular infection and with proteasome blockade treatment, we also found that *pals-5*p::GFP reporter expression was much lower or absent in *zip-1(jy14)* mutants than in wild-type animals, and indistinguishable from the phenotype of *zip-1(jy13)* mutants (Fig. S3). In summary, we found that phenotypes observed after loss of *zip-1* are not allele-specific and that they likely cannot be attributed to inactivation of the *y75b8a.55* gene. Altogether, these results indicate that *zip-1* controls expression of *pals-5* reporters induced by all well-characterized triggers of IPR gene expression.

### *zip-1* is required for early induction of *pals-5* mRNA as well as induction of a subset of other IPR genes

We next used qRT-PCR to assess the role of *zip-1* in controlling levels of endogenous *pals-5* mRNA, as well as mRNA of other IPR genes. Because bortezomib treatment induced the strongest and most consistent IPR gene expression, we used this trigger to assess the role of *zip-1* in mediating IPR gene induction in subsequent experiments. Here we were surprised that *zip-1(jy13)* mutants had only about a 3.5 fold reduction in *pals-5* mRNA induction at 4 hours (h) after bortezomib treatment compared to induction in wild-type animals (Fig. 3A). 4 h is the timepoint at which *zip-1* mutants were strongly defective for induction of *pals-5*p::GFP and *pals-5*p::NanoLuc expression (Fig. 2E, F, G). Therefore, we considered the possibility that GFP and nanoluciferase expression observed at 4 h may reflect protein synthesized from mRNA made earlier. To investigate this possibility, we used qRT-PCR to measure *pals-5* mRNA at 30 minutes (min) after bortezomib treatment, and here we found that *zip-1* was completely required for the ∼300-fold induction of *pals-5* mRNA at this early timepoint (Fig. 3B). Thus, *zip-1* is completely required for *pals-5* mRNA induction 30 min after bortezomib treatment, but only partially required for induction at 4 h after bortezomib treatment.

**Fig. 3.**
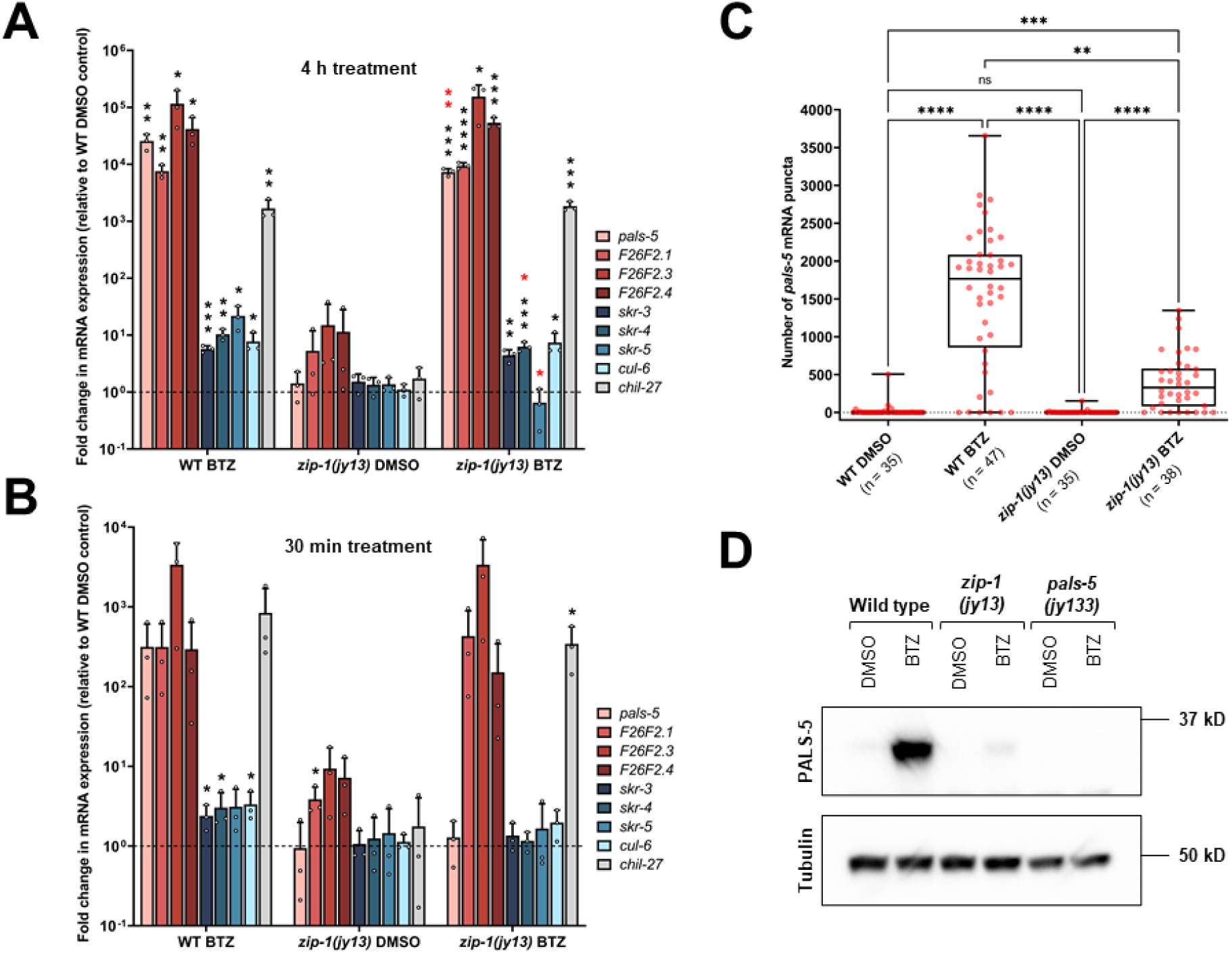
*zip-1* regulates the early phase of *pals-5* transcription following bortezomib treatment, and controls some IPR gene expression. (A, B) qRT-PCR measurements of selected IPR genes and *chil-27* at 4 h timepoint (A) and 30 min timepoint (B) of bortezomib (BTZ) or DMSO treatments. The results are shown as the fold change in gene expression relative to wild-type DMSO diluent control. Three independent experimental replicates were analyzed; the values for each replicate are indicated with circles. Error bars represent standard deviations. A one-tailed t-test was used to calculate *p*-values; black asterisks represent significant difference between the labeled sample and the wild-type DMSO control; red asterisks represent significant difference between wild-type (WT) N2 and *zip-1(jy13)* bortezomib treated samples; **** *p* < 0.0001; *** *p* < 0.001; ** *p* < 0.01; * 0.01 < *p* < 0.05; *p*-values higher than 0.05 are not labeled. (C) smFISH quantification of number of *pals-5* mRNA transcripts in the first four intestinal cells. Three experimental replicates were performed and at least 33 animals were analyzed for each sample (at least five animals were analyzed per sample per replicate), 4 h after bortezomib or DMSO control treatment. In box-and-whisker plots, each box represents 50% of the data closest to the median value (line in the box), whereas whiskers span the values outside of the box. A Kruskal-Wallis test was used to calculate *p*-values; **** *p* < 0.0001; *** *p* < 0.001; ** *p* < 0.01; ns indicates nonsignificant difference (*p* > 0.05). (D) Western blot analysis of PALS-5 expression in wild-type, *zip-1(jy13)* and *pals-5(jy133)* animals. *pals-5(jy133)* is a complete deletion of the *pals-5* gene and was used as a negative control. Animals were treated with bortezomib or DMSO control for 4 h. PALS-5 was detected using anti-PALS-5 antibody, whereas anti-tubulin antibody was used as a loading control. Predicted sizes are 35.4 kD for PALS-5 and around 50 kD for different members of tubulin family.

Because *zip-1* appeared to be more important at 4 h for inducing GFP and nanoluciferase transcriptional reporters than for inducing *pals-5* mRNA by qRT-PCR, we used smFISH as a separate measure for *pals-5* mRNA levels at this timepoint. Here, as in the GFP reporter studies, *pals-5* expression was seen in the intestine. Because it is an easily identified location, we quantified *pals-5* RNA levels in the first intestinal ring, which is comprised of four epithelial cells. Here we found that *pals-5* mRNA was induced to a lesser degree in *zip-1* mutants treated with bortezomib compared to wild-type animals (Fig. 3C, Fig. S4).

Next, to determine whether *zip-1* mutants are defective in PALS-5 protein production, we raised polyclonal antibodies against the PALS-5 protein. Using these antibodies for western blots, we found that PALS-5 protein induction in *zip-1(jy13)* animals at 4 h after bortezomib treatment was almost undetectable in comparison to the induction in bortezomib-treated wild-type animals (Fig. 3D, Fig. S5). Therefore, *zip-1* is required for high levels of PALS-5 protein production after bortezomib treatment, very likely through its role in regulating induction of *pals-5* mRNA.

Having confirmed that *zip-1* is completely required for induction of *pals-5* mRNA at 30 min and partially required at 4 h, we examined the requirement for *zip-1* in induction of other IPR genes at these timepoints. We analyzed highly induced IPR genes of unknown function – *F26F2.1*, *F26F2.3* and *F26F2.4*, as well as components of a cullin-ring ubiquitin ligase complex – *cul-6*, *skr-3*, *skr-4* and *skr-5,* which mediates thermotolerance as part of the IPR program (18). Interestingly, *zip-1* was not required at either time point (30 min or 4 h after bortezomib treatment) for mRNA induction of the majority of genes we analyzed, including *F26F2.1* (Fig. 3 A and B). In agreement with these results, *zip-1* was not required for *F26F2.1*p::GFP expression after bortezomib treatment (Fig. S6). Furthermore, *zip-1* was not required for induction of the chitinase-like gene *chil-27*, which is induced by bortezomib, as well as by the natural oomycete pathogen *Myzocytiopsis humicola* (13, 23). In contrast, *zip-1* was required at the 4 h timepoint for induction of *skr-5* RNA levels (Fig. 3B). Because the induction of *skr-5* at 30 min was quite low, it was difficult to assess the role of *zip-1* in regulating this gene at this timepoint. Overall, these results suggest that there are at least three classes of IPR genes: 1) genes that require *zip-1* for early but not later induction (“Early *zip-1*-dependent” genes like *pals-5*), 2) genes that require *zip-1* at the later timepoint (“Late *zip-1*-dependent” genes like *skr-5*), and 3) genes that do not require *zip-1* at either timepoint for their induction (“*zip-1*-independent” genes like *F26F2.1*).

We also analyzed the role of *zip-1* in regulating IPR gene induction upon intracellular infection. Similar to the results using bortezomib as a trigger, we found that *zip-1* was required for *pals-5* induction by *N. parisii* infection or by Orsay virus infection, but was not required for induction of F26F2.1. Interestingly, *zip-1* mutation had only a partial effect on *skr-5* induction following Orsay-virus infection, whereas *skr-5* levels were not highly induced in *N. parisii* infected animals (Fig. S7).

### RNA sequencing analysis reveals a genome-wide picture of *zip-1*-dependent genes

To obtain a genome-wide picture of the genes controlled by *zip-1*, we next performed RNA sequencing (RNA-seq) analysis. Here we treated wild-type N2 or *zip-1(jy13)* mutant animals with either bortezomib or vehicle control for either 30 min or 4 h, then collected RNA and performed RNA-seq. Based on differential expression analyses, we created lists of genes upregulated in each genetic background after bortezomib treatment at both analyzed timepoints. At 30 min, we found that 136 and 215 genes were upregulated in wild-type and *zip-1(jy13)* animals, respectively, with 72 genes being upregulated in both backgrounds (Fig. 4A, Table S2). Therefore, 64 genes (i.e. 136 minus 72 genes) were induced only in wild-type animals, indicating that they are *zip-1*-dependent early upon proteasome blockade. Importantly, *pals-5* was among these genes that were only upregulated in wild-type animals and not *zip-1* mutants at this timepoint, consistent with our qRT-PCR analysis (Fig. 3). At 4 h, we identified many more genes that showed differential expression between bortezomib and control treatments in both genetic backgrounds, with 2923 and 2813 genes upregulated in wild-type and *zip-1(jy13)* mutants, respectively (Fig. 4B, Table S2). 2035 genes were upregulated in both backgrounds, meaning that 888 genes (2923 minus 2035 genes) were specifically upregulated in wild-type animals. 883 out of 888 genes belong to the “Late *zip-1*-dependent” category, and include *skr-5*, consistent with our qRT-PCR analysis (Fig. 3). Notably, five genes (*ZK355.8*, *K02E7.10*, *math-39*, *gst-33* and *F55G1.7*) were induced only in wild-type animals at both examined timepoints, and thus we classified these genes as “Completely *zip-1*-dependent”. Therefore, 59 (64 minus 5) genes from the 30 min timepoint belong to the “Early *zip-1*-dependent” genes category. Of note, consistent with our qRT-PCR and GFP reporter analysis, the *F26F2.1* gene was upregulated in both genetic backgrounds following bortezomib treatment, and thus belongs to the “*zip-1*-independent” category.

**Fig. 4.**
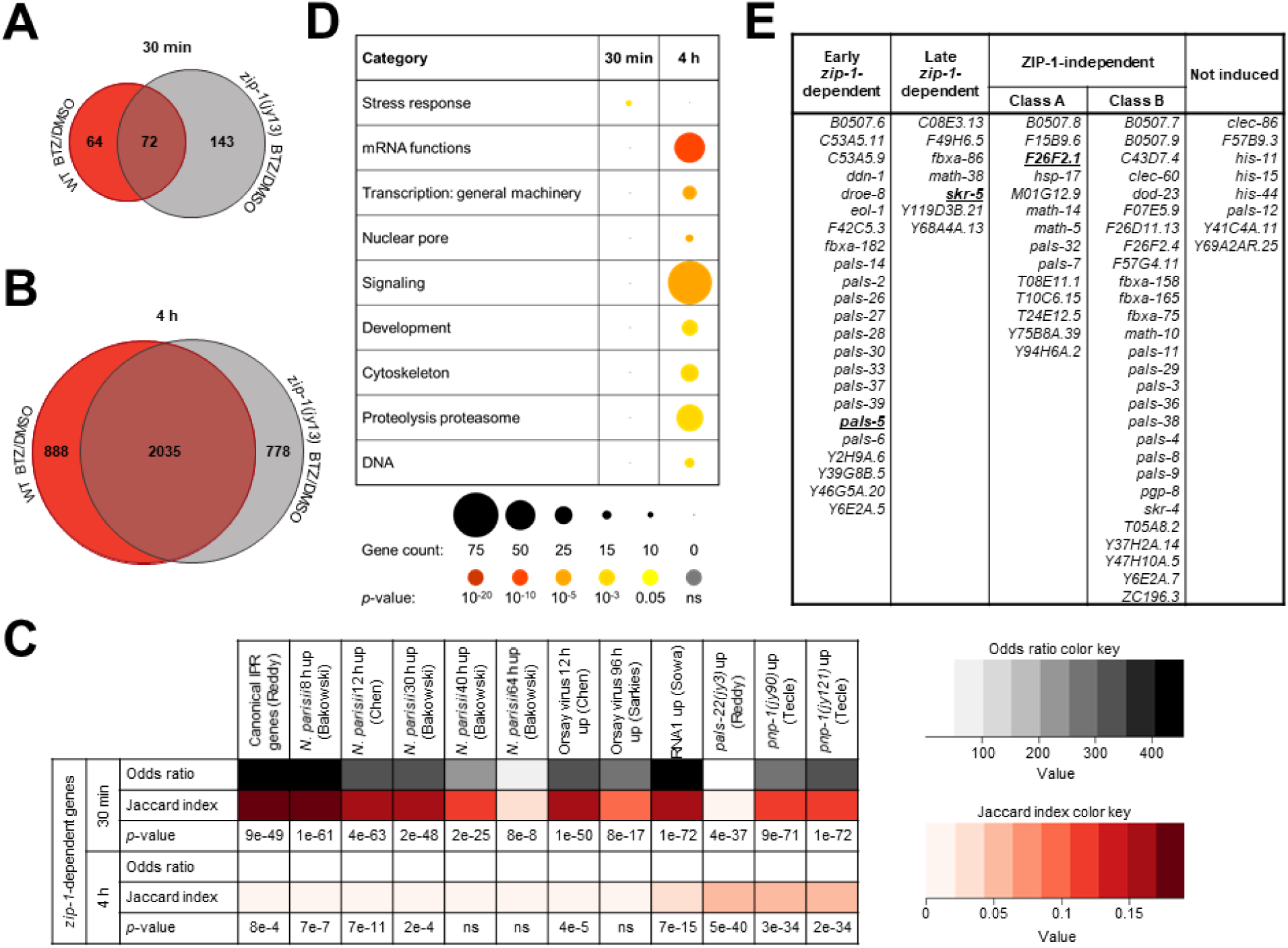
Defining *zip-1*-dependent IPR genes. (A, B) Venn diagrams of differentially expressed genes following 30 min (A) and 4 h bortezomib treatments (B) in WT N2 and *zip-1(jy13)* mutant animals as compared to DMSO controls for each background. 64 and 888 genes were upregulated after 30 min and 4 h bortezomib treatment in N2 animals, respectively, but not in *zip-1(jy13)* mutants, suggesting that these genes are *zip-1*-dependent. (C) The list of *zip-1*-dependent genes shows significant overlap with previously published list of genes that are upregulated by different IPR triggers. A Fisher’s exact test was used to calculate odds ratios and *p*-values. These values were calculated taking in account all genes in *C. elegans* genome. If the odds ratio is greater than one, two data sets are positively correlated. Jaccard index measures similarity between two sets, with the range 0-1 (0 – no similarity, 1 – same datasets). For approximate quantification, the odds ratio and Jaccard index color keys are indicated on the right side of the table. (D) Graphical representation of enriched gene categories for all *zip-1*-dependent genes at 30 min and 4 h timepoints of bortezomib treatment. Each category represents a biological process or a structure associated with *zip-1*-dependent genes at either timepoint. Count of genes found in each category is indicated by the circle size, as illustrated under the table. Statistical significance for each category is indicated by the circle color; *p*-values are indicated under the table. (E) Classification of 80 canonical IPR genes based on their ZIP-1 dependency. Representative canonical IPR genes from each class are shown in bold and underlined.

We next examined the correlation between *zip-1*-dependent genes (separately analyzing genes induced at each timepoint) and gene sets that were previously associated with IPR activation. Here we found that there is a significant similarity between *zip-1*-dependent genes induced after 30 min bortezomib treatment, and genes upregulated early after Orsay virus infection, *N. parisii* infection, ectopic expression of Orsay viral RNA1, and genes induced in *pals-22* and *pnp-1* mutants (Fig. 4C, Table S3). In addition, there is a significant similarity between genes induced at 30 min timepoint and canonical IPR genes. Similarly, there is a significant overlap between *zip-1*-dependent genes induced after 4 h bortezomib treatment and the majority of these IPR-associated gene-sets. Of note, there was not a significant overlap between *zip-1*-dependent genes induced after 4 h bortezomib treatment, and genes that are upregulated at the late phases of viral (96 hpi) and microsporidia infections (40 hpi and 60 hpi). These results suggest that *zip-1* plays a more important role in the acute transcriptional response to intracellular infection, and perhaps a lesser role later in infection. Furthermore, our analysis revealed significant similarity between *zip-1*-upregulated genes and genes that are downregulated by *sta-1*. STA-1 is a STAT-related transcription factor that acts as a negative regulator of IPR gene expression. (Fig. S8A, Table S3). We also found a significant overlap between *zip-1*-dependent genes and those induced by *M. humicola*, a natural oomycete pathogen that infects the epidermis, although *zip-1* was not required for induction of the chitinase-like gene *chil-27*, which is a common marker for *M. humicola* response (Fig. S8B, Table S3). Previous studies have shown connections between the IPR and genes induced either by *M. humicola* infection, or by extract from *M. humicola* as part of the oomycete recognition response in the epidermis (13, 23, 24).

We identified *zip-1*-dependent genes in our analysis here using proteasome blockade by bortezomib, which has effects on transcription that are unrelated to the IPR. For example, bortezomib activates the bounceback response that induces expression of proteasome subunits, and it is controlled by the conserved transcription factor SKN-1/Nrf2 (25). Therefore, we compared if *zip-1*-dependent genes (from both analyzed timepoints) have significant overlap with *skn-1*-dependent genes. Here we found no significant similarity between the majority of analyzed datasets (Fig. S8C, Table S3). These results are consistent with previous IPR RNA-seq studies showing a distinction between the IPR and the bounceback response, and suggest that *zip-1* does not play a role in the bounceback response (11, 13).

In addition, we found that *zip-1* mRNA itself was strongly upregulated in wild-type animals following bortezomib treatment, consistent with previous studies (13). Surprisingly however, we found that *zip-1* mRNA was also upregulated in *zip-1* mutants. This result that was initially confusing, because the *zip-1* coding sequence is completely deleted in the *zip-1(jy13)* allele that we used in RNA-seq analysis. Upon closer examination however, we found that *zip-1* sequencing reads in *zip-1(jy13)* mutant samples aligned to the region upstream of the *zip-1* gene coding sequence, which contains annotated 5’ untranslated regions (UTRs) for several *zip-1* isoforms, as well as to downstream sequences that contain the *zip-1* 3’ UTR (Fig. S9). This finding indicates that *zip-1* is not required to induce its own transcription, but rather a distinct transcription factor is involved in upregulation of *zip-1* mRNA expression.

To obtain insight into other biological processes and cellular structures that may be related to *zip-1*, we performed analysis with the WormCat program, specifically designed for analysis of *C. elegans* genomics data (26). We separately analyzed 64 genes from the early timepoint and 888 genes from the later timepoint that were upregulated in wild-type animals but not *zip-1* mutants. The only significantly overrepresented category of upregulated genes at 30 min was the stress response category (Fig. 4D, Table S4). Analysis of the genes upregulated at 4 h revealed a significant overrepresentation of genes implicated in mRNA function, transcription, nuclear pore, signaling, development, cytoskeleton, proteolysis and DNA.

Finally, we analyzed and classified 80 canonical IPR genes (13) based on their expression levels in our RNA-seq datasets, to place them into different categories based on their dependence on *zip-1*. Here we found that 23 IPR genes (including *pals-5*) were upregulated in wild-type animals but not *zip-1* mutants 30 min after bortezomib treatment, but became upregulated in both genetic backgrounds at 4 h (Fig. 4E). Therefore, these genes are “Early *zip-1*-dependent” IPR genes. Notably, 11 *pals* genes belong to this category. Another seven IPR genes (including *skr-5*) were not upregulated in *zip-1(jy13)* mutants at either timepoint analyzed, but were upregulated in wild type at 4 h, and we classified these genes as “Late *zip-1*-dependent” IPR genes. Therefore, overall, 30 IPR genes appeared to be *zip-1*-dependent, when including both timepoints. 42 canonical IPR genes were upregulated in both genetic backgrounds, and we classified them as “*zip-1*-independent” IPR genes. Because some of these genes were not upregulated at the first timepoint, we further divided this category of genes into class A that showed upregulation after 30 min bortezomib treatment (including *F26F2.1*), and class B that showed upregulation only after 4 h of bortezomib treatment. Of note, eight canonical IPR genes did not show significant upregulation after bortezomib treatment, so we did not classify them in any category. These include histone genes, which previous studies had shown to be regulated by *pals-22/pals-25* and *N. parisii* infection (and thus qualify as IPR genes), but not to be induced by bortezomib treatment (11, 13). In conclusion, our RNA-seq results demonstrate that *zip-1* controls RNA expression of 30 out of 80 IPR genes, and reveal that IPR genes can be placed into three separate classes based on their regulation by *zip-1*.

### ZIP-1 is expressed in the intestine and is required in this tissue to regulate *pals-5* gene expression

To examine where ZIP-1 is expressed, we tagged the *zip-1* endogenous genomic locus with *gfp* immediately before the stop codon using CRISPR/Cas9-mediated gene editing. Here we found that ZIP-1::GFP endogenous expression was not detectable in unstressed animals. Because *zip-1* mRNA is induced by bortezomib, and bortezomib blocks protein degradation, we investigated whether ZIP-1::GFP was visible after bortezomib treatment. Here we found that ZIP-1::GFP expression was induced, with strongest expression found in intestinal nuclei (Fig. 5A). Nuclear expression was also identified in the epidermis (Fig. 5B). Specifically, 98 % (59/60) of animals showed ZIP-1::GFP expression in intestinal nuclei after 4 h bortezomib treatment, while 88 % (53/60) showed expression in epidermal nuclei after 4 h bortezomib treatment. In contrast, no GFP signal was observed in wild-type animals treated with bortezomib, or in *zip-1::gfp* mutants or wild-type animals treated with DMSO control (60 analyzed animals for each condition).

**Fig. 5.**
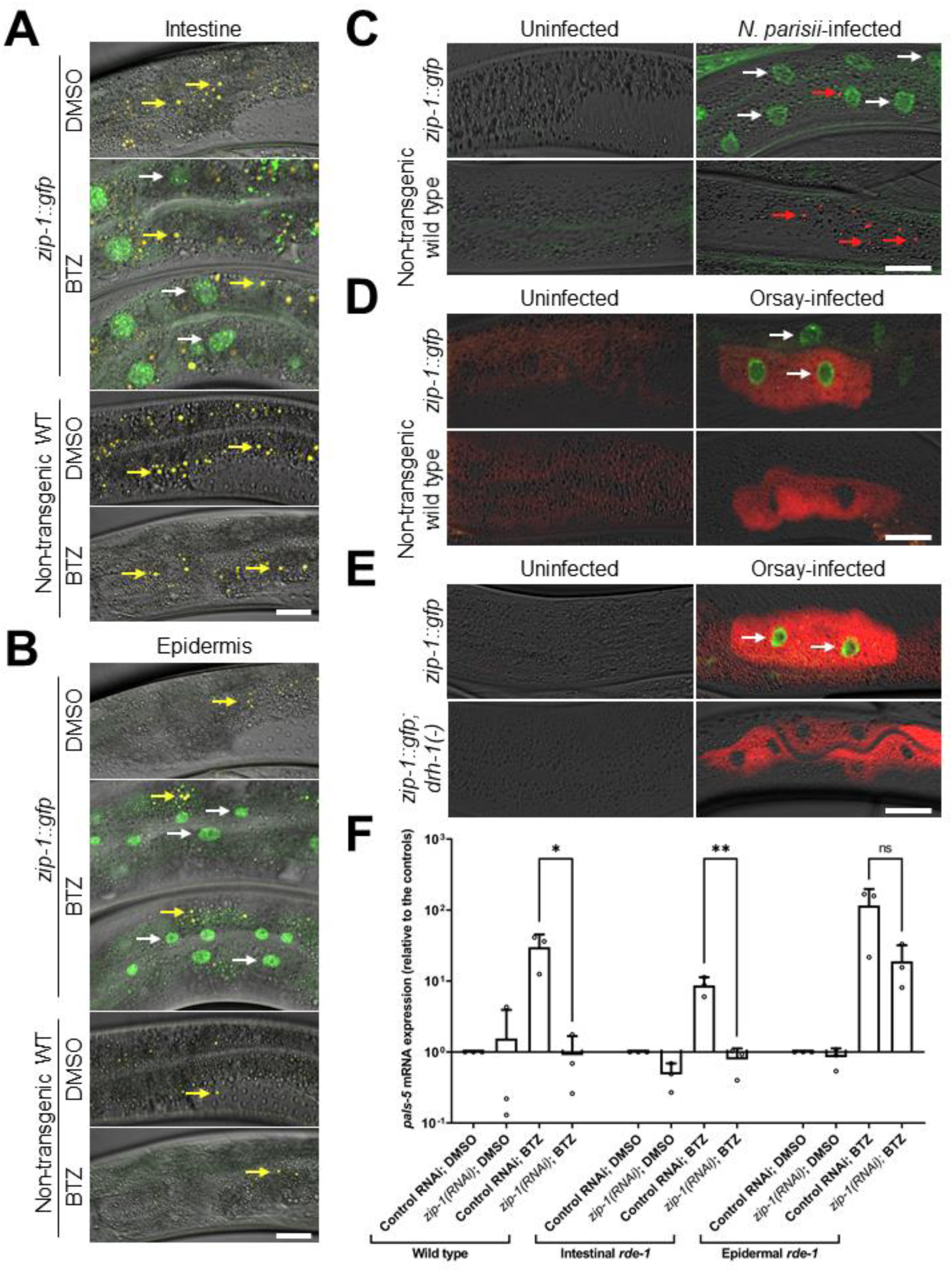
*zip-1* acts in the intestine to regulate *pals-5* mRNA levels. (A, B) ZIP-1::GFP is expressed in intestinal (A) and epidermal nuclei (B) 4 h after bortezomib treatment. No expression was observed in animals exposed to DMSO control, or in the non-transgenic control strain N2. Composite images consist of merged fluorescent (GFP and autofluorescence) and DIC channels. Yellow signal in the composite images depicts autofluorescence from gut granules. White arrows indicate representative intestinal nuclei expressing ZIP-1::GFP; yellow arrows indicate autofluorescence. Scale bar = 20 µm. (C-E) *N. parisii* (C) and Orsay virus infection (D, E) induce ZIP-1::GFP expression in intestinal nuclei. Fluorescent and DIC images were merged; green represents ZIP-1::GFP; fluorescence from pathogen-specific FISH probes is shown in red. White arrows indicate representative intestinal nuclei expressing ZIP-1::GFP (C-E); red arrows indicate *N. parisii* sporoplasms (C). Scale bar = 30 µm. (F) Intestine-specific *zip-1(RNAi)* prevents *pals-5* mRNA induction. qRT-PCR measurements of *pals-5* levels at the 30 min timepoint of bortezomib (BTZ) or DMSO treatments. The results are shown as fold change in gene expression relative to DMSO diluent control. Three independent experimental replicates were analyzed; the values for each replicate are indicated with circles. Error bars represent standard deviations. A one-tailed t-test was used to calculate *p*-values; ** *p* < 0.01 is indicated with two asterisks; * 0.01 < *p* < 0.05; ns indicates nonsignificant difference (*p* > 0.05).

Next we examined ZIP-1::GFP expression upon intracellular infection. First, we infected ZIP-1::GFP animals with *N. parisii*, stained with a FISH probe to label parasite cells, and quantified the percentage of infected animals displaying ZIP-1::GFP nuclear expression. Here we found that 73.33% (44/60) of animals with parasite cells had ZIP-1::GFP expression, whereas 0% (0/60) of uninfected animals had ZIP-1::GFP expression (Fig. 5C). We did note that uninfected intestinal cells found adjacent to *N. parisii*-infected cells also displayed ZIP-1::GFP expression, suggesting there may be cell-to-cell signaling from infected to uninfected cells to induce ZIP-1::GFP expression, although we cannot eliminate the possibility that ‘uninfected’ cells have pathogen that was not visible. Next we infected ZIP-1::GFP animals with Orsay virus, and stained with a FISH probe to label viral RNA. Similar to studies with *N. parisii*, we found that 100% (60/60) virus infected animals had ZIP-1::GFP expression, whereas 0% (0/60) of uninfected animals had ZIP-1::GFP expression (Fig. 5D). Here as well, we found evidence there may be cell-to-cell signaling, as uninfected cells adjacent to viral-infected cells displayed ZIP-1::GFP expression. Importantly, viral induction of ZIP-1::GFP enabled us to analyze whether the DRH-1 acts upstream of ZIP-1::GFP. Here we found that viral infection no longer induced ZIP-1::GFP in *drh-1* mutants (0% or 0/60 animals) (Fig. 5E), suggesting that the DRH-1 receptor acts upstream of the ZIP-1 transcription factor.

To determine the tissue in which *zip-1* acts to regulate *pals-5* induction, we performed tissue-specific downregulation of *zip-1* using RNAi, and measured the levels of *pals-5* mRNA following 30 min bortezomib treatment. First, we used *rde-1* loss-of-function mutation strains, which have a *rde-1* rescuing construct expressed specifically in either the intestine or in the epidermis, which leads to enrichment of RNAi in these tissues. Here we observed that *zip-1* RNAi in the intestinal-specific RNAi strain caused a block in *pals-5* induction, similar to *zip-1* RNAi in wild-type animals (Fig. 5F). In contrast, *pals-5* induction was less compromised by *zip-1(RNAi)* in the epidermal-specific RNAi strain. Because intestinal expression of *rde-1* allows generation of secondary siRNAs that can spread to other tissues and silence gene expression there, we used a separate tissue-specific RNAi strain, where the *sid-1* transport channel is specifically expressed in the intestine (27, 28). These mutants do not suffer from the problem of leakiness seen in the *rde-1* rescue strains (28, 29). However, we did note that they suffer from the opposite problem: they appear to be somewhat resistant to RNAi. To quantify this effect, we treated the intestinally rescued *sid-1* strain with dsRNA against *act-5*, which is an actin isoform expressed in the intestine, but not in the epidermis, and it is essential for development (30). Here we found that *act-5* RNAi caused less severe effects on size in the intestinally rescued *sid-1* strain compared to wild-type animals (Fig. S10A and S10B). Despite being partially resistant to RNAi, we did find that *zip-1* RNAi in this strain caused a significant reduction in *pals-5*mRNA induction upon bortezomib treatment compared to vector control (Fig. S10C). Taken together, our data suggest that *zip-1* is highly expressed in the intestinal nuclei following bortezomib treatment, and that *zip-1* is important in the intestine for induction of *pals-5* mRNA.

### *zip-1* regulates resistance to natural intracellular pathogens

Because increased IPR gene expression is correlated with increased resistance to intracellular infection (10, 13), we investigated the role of *zip-1* in resistance to intracellular pathogens. First, we investigated Orsay virus. Here, we infected L4 animals and found that *zip-1* mutants had higher viral load compared to wild-type animals, as assessed by qRT-PCR (Fig. 6A). Similarly, we found upon infection of L1 animals and measuring viral load with FISH staining that *zip-1* mutants had a trend toward higher infection rate than wild-type animals (Fig. 6B). We also investigated whether *zip-1* might have a greater effect on viral load in a mutant background where IPR genes are constitutively expressed. Indeed, we found a more pronounced role for *zip-1* after viral infection of *pnp-1* mutants, which have constitutive expression of IPR genes, including *pals-5* (Fig. 6B) (15). Of note, our qRT-PCR analysis of *pnp-1(jy90)* animals showed that elevated *pals-5* mRNA levels depend on *zip-1*, suggesting that the IPR genes upregulated by *zip-1* promote resistance against viral infection (Fig. S11). Similar to what we observed after bortezomib treatment, expression of highly induced IPR genes *F26F2. 1*, *F26F2.3* and *F26F2.4* in a *pnp-1* mutant background did not require *zip-1*. This finding suggests that *zip-1-*dependent IPR genes may play a more important role in Orsay virus resistance than other IPR genes.

**Fig. 6.**
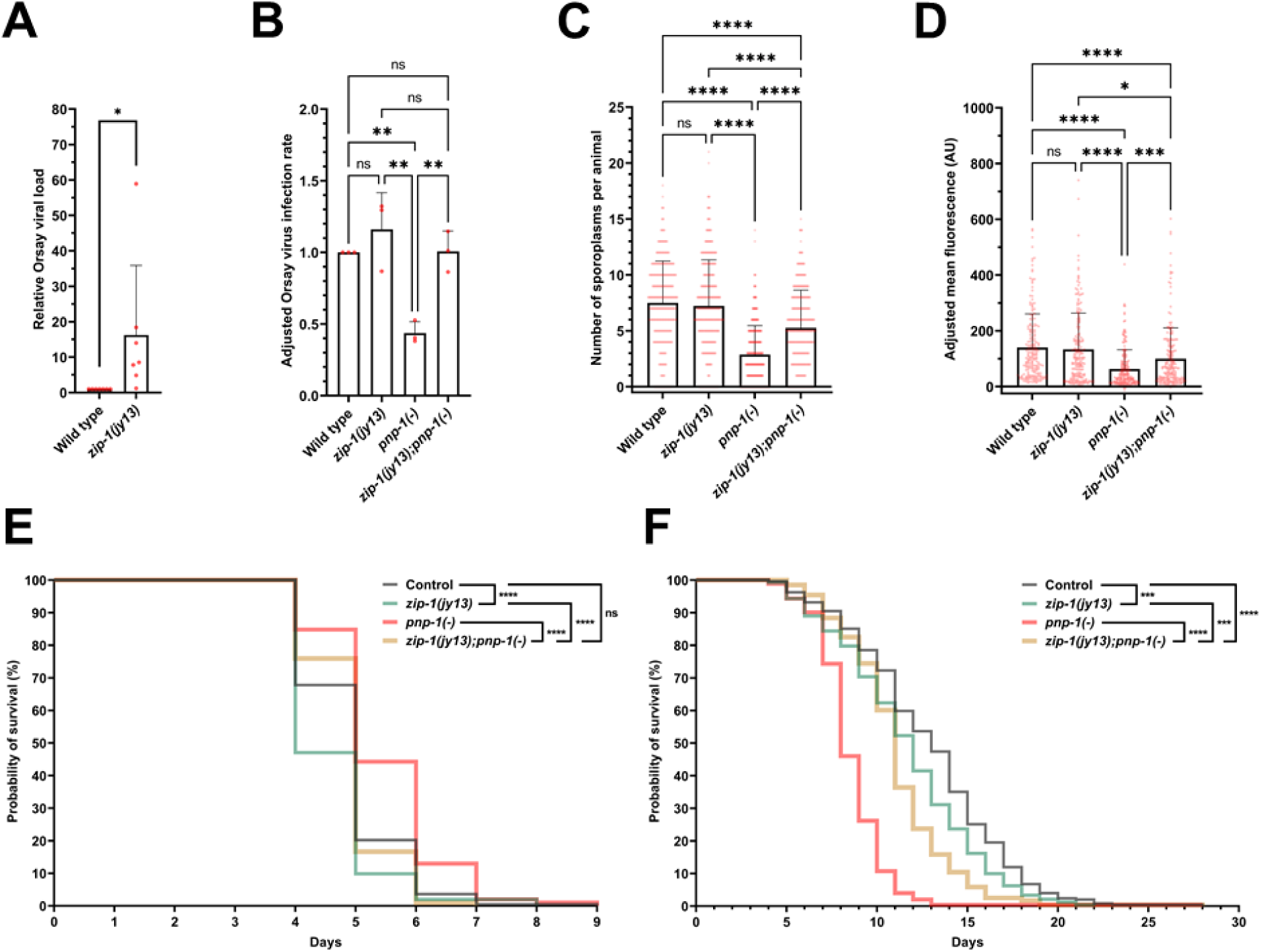
*zip-1* promotes resistance to intracellular pathogens. (A) qRT-PCR analysis of Orsay virus RNA1 levels in control and *zip-1(jy13)* mutant animals. Animals were infected at L4 stage and collected at 24 hpi. Four experimental replicates were analyzed, each consisting of two biological replicates assayed in technical duplicates. (B) Fraction of animals infected with Orsay virus in control, *zip-1(jy13)*, *pnp-1(jy90)* and *zip-1(jy13); pnp-1(jy90)* backgrounds at 12 hpi. Animals were infected at L1 stage. 900 animals per strain were scored based on the presence or absence of the Orsay virus RNA1-specific FISH probe fluorescence. The infection rate of the control strain was set to one. (C) *N. parisii* pathogen load quantified at 3 hpi as number of sporoplasms per animal; 300 L1 animals were analyzed per strain in three experimental replicates. (D) Quantification of *N. parisii*-specific mean FISH fluorescence signal normalized to body area excluding pharynx. Animals were infected at L1 stage and analyzed at 30 hpi; 200 animals were analyzed per strain. The head region was excluded from the analysis because of the expression of the red coinjection marker *myo-2*p::mCherry. In box-and-whisker plots, each box represents 50% of the data closest to the median value (line in the box), whereas whiskers span the values outside of the box. AU – arbitrary units. (A-D) All strains are in a *pals-5p::gfp* background. (E) Survival of wild-type, *zip-1(jy13)*, *pnp-1*(*-*) and *zip-1(jy13); pnp-1*(*-*) animals following *N. parisii* infection. Animals were exposed to *N. parisii* spores for 66 h from L1 stage, and then transferred daily to non-infectious plates and scored for viability. (F) Longevity analysis of strains used in the infection assays. (E, F) Animals were incubated at 25°C. Data from 3 experimental replicates are shown in a single graph. Percentage of alive animals is indicated on y axis for each day of analysis (x axis). Statistical analyses were performed using an unpaired t-test (A), an ordinary one-way ANOVA (B), a Kruskal-Wallis (C, D) and a log-rank (Mantel-Cox) test (E, F) to calculate *p*-values; **** *p* < 0.0001; *** *p* < 0.001; ** *p* < 0.01; * 0.01 < *p* < 0.05; ns indicates nonsignificant difference (*p* > 0.05).

Next, we examined a role for *zip-1* in resistance to *N. parisii* infection by measuring pathogen load. Here we did not see an effect of *zip-1* in a wild-type background either at 3 hpi or at 30 hpi (Fig. 6 C and D). However, at both timepoints, we found that loss of *zip-1* significantly suppressed the increased pathogen resistance (i.e. lower pathogen load) of *pnp-1* mutants (Fig. 6 C and D). Therefore, these experiments indicate that wild-type *zip-1* promotes resistance to *N. parisii* infection in a background where IPR genes are induced prior to infection. To further analyze the role of *zip-1* in response to *N. parisii* infection, we also performed killing assays in which we analyzed survival of animals following infection. Consistent with published data, we found that *pnp-1* mutants survive longer than wild-type animals when infected with *N. parisii*, but do not survive longer than wild-type animals in the absence of infection (Fig. 6 E and F). Importantly, we found that *zip-1* mutations decrease survival both in a *pnp-1* mutant background, as well as in a wild-type background. Therefore, wild-type *zip-1* promotes survival against *N. parisii* infection. Because infections were performed by feeding pathogens to animals, it was possible that differences in food intake and elimination were responsible for any differences seen in pathogen load. Therefore, we measured accumulation of fluorescent beads in all tested strains and we did not find any significant differences between *zip-1* mutants and control animals (Fig. S12). In conclusion, the increased pathogen load in *zip-1* mutants is unlikely to be due to differences in the exposure of intestinal cells to pathogen in these mutants.

Other phenotypes in *pnp-1* mutants include higher sensitivity to heat shock and slightly slower growth rate (15). We tested if either of these phenotypes are *zip-1*-dependent. First, we found that *zip-1(jy13)* animals had a similar survival rate after heat shock compared to the control strain (Fig. S13A). Similarly, we found that loss of *zip-1* in a *pnp-1(jy90)* mutant background did not significantly suppress the higher lethality observed in *pnp-1(jy90)* single mutants, suggesting that ZIP-1 does not play a crucial role in thermotolerance regulation. Finally, we analyzed if *zip-1(jy13)* mutants, which show a wild-type growth rate, can suppress the mild growth delay caused by a *pnp-1(jy90)* mutation. Here, growth was assayed based on the body length measurements 44 h after plating synchronized L1 animals, and we found that *zip-1(jy13); pnp-1(jy90)* animals were still significantly smaller than control animals and *zip-1(jy13)* single mutants (Fig. S13B). Therefore, *zip-1* does not appear to be important for these non-infection related phenotypes of *pnp-1* mutants. Instead, it seems that *zip-1* specifically plays a role in regulating immunity-related IPR genes.

## Discussion

Most studies of antiviral immunity in invertebrates have focused on anti-viral RNAi, and less is known about transcriptional responses to intracellular infection in either of the two major invertebrate model systems, *Drosophila melanogaster* or *C. elegans* (31–33). The IPR in *C. elegans* is a common transcriptional response that is induced independently by both virus and microsporidia infection, as well as by specific physiological perturbations such as proteotoxic stress (11–13). Previous studies had shown that the STAT-related transcription factor *sta-1* was a repressor of IPR genes (34), but the activating transcription factor for the IPR was not known. Here, we show that the previously uncharacterized, predicted bZIP transcription factor ZIP-1 functions downstream of all known IPR triggers to induce a subset of IPR genes (Fig. 7). Importantly, we show that *zip-1* plays a role in immunity against infection by both the Orsay virus and microsporidia. Therefore, *zip-1* appears to be the first transcription factor shown to promote an inducible defense response against intracellular intestinal pathogens in *C. elegans*.

**Fig. 7.**
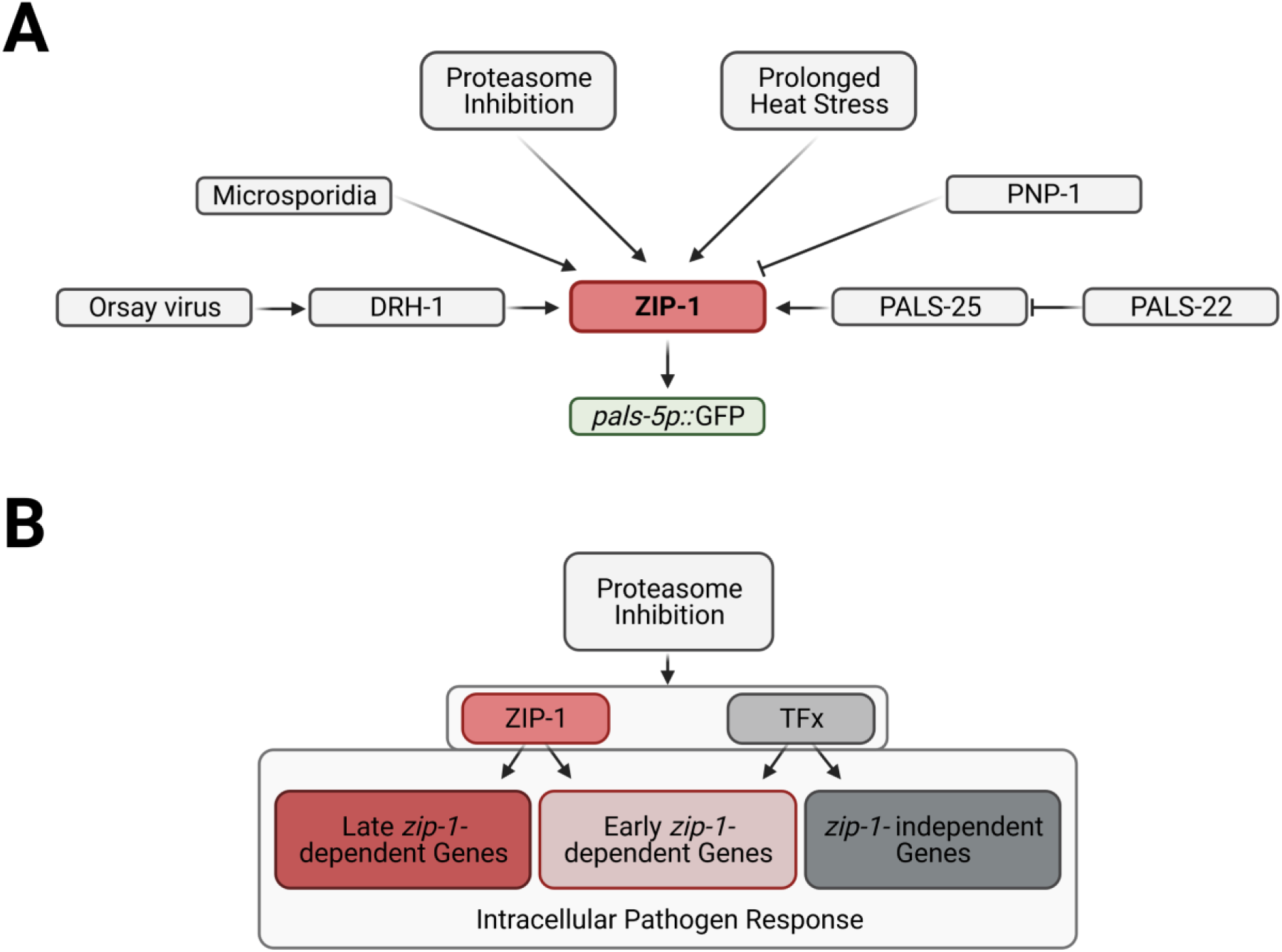
Model of IPR gene regulation. (A) All known IPR activating pathways require ZIP-1 for induction of the *pals-5*p::GFP reporter. (B) IPR genes can be divided into three categories: early *zip-1*-dependent, late *zip-1*-dependent and *zip-1*-independent genes. Unknown transcription factor or factors (TFx) regulate expression of early *zip-1*-dependent genes at later timepoint, as well as transcription of *zip-1*-independent genes.

ZIP-1 adds to the growing list of bZIP transcription factors involved in *C. elegans* immunity. The bZIP transcription factor family is expanded in *C. elegans* compared to other organisms (35) and several members of this family have been previously implicated in defense against the extracellular bacterial pathogens of the intestine. For example, the central pathway in defense against bacterial pathogens like *Pseudomonas aeruginosa* is the p38 MAPK pathway, which leads to activation of the ATF-7 bZIP transcription factor, as well as the bZIP-related transcription factor SKN-1 in response to reactive oxygen species generated upon infection (36, 37). The bZIP proteins ZIP-2 and CEBP-2 control induction of several p38-independent genes induced by *P. aeruginosa*, in response to the *P. aeruginosa* translation-blocking ExotoxinA (38–41). Under certain infection conditions, ZIP-2 and CEBP-2 act with two other bZIP transcription factors, ZIP-4 and CEBP-1, to control induction of the Ethanol and Stress Response Element network upon *P. aeruginosa* infection, likely in response to mitochondrial damage (42). Furthermore, the bZIP proteins ATFS-1 and ZIP-3 have been shown to play antagonistic roles in activation of mitochondrial unfolded protein response upon damage caused by *P. aeruginosa* infection (43, 44).

In addition to the bZIP transcription factors mentioned above, several other classes of transcription factors play roles in *C. elegans* defense, including FOXO, GATA, HSF, HLH and NHR transcription factor family members (45–50). Moreover, several members of *C. elegans* Myc family of transcription factors have been shown to be the regulators of microsporidia growth and development (19). What is the logic to having so many transcription factors involved in immunity in *C. elegans*? For comparison, only one bZIP transcription factor, CrebA, has recently been shown to play a role in *D. melanogaster* tolerance to bacterial pathogens (51). Also, a single STAT transcription factor – a component of JAK/STAT pathway, has been shown to play a downstream role in antiviral immunity, although this factor is not thought to be the first responder to viral infection (52, 53). The majority of studies in *D. melanogaster* indicate that the NF-kB transcription factors Dif, Dorsal and Relish are the major transcription factors to induce immune genes upon bacterial and fungal infections, and they also play a role in antiviral immunity (53–55). A large percentage of immune genes in humans are also controlled by NF-kB transcription factors upon induction by bacterial infection, and by IRF3/7 upon viral infection, working together with NF-kB (56–59). Interestingly, NF-kB was lost in the evolutionary lineage that gave rise to *C. elegans*, so perhaps several other transcription factors fill that gap to induce defense (14). Or perhaps, this diverse list of immune-related transcription factors is a result of *C. elegans* apparently lacking professional immune cells like macrophages or hemocytes, which play key roles in mammalian and *D. melanogaster* defense respectively (60, 61). For this reason, studies in *C. elegans* have focused on non-professional immune cells like epithelial cells (62–64), which are less well studied in mammalian research compared to professional immune cells like macrophages. If more mammalian and *D. melanogaster* studies screened for transcription factors acting in epithelial cells, the lists might grow longer there as well.

Although our study indicates that ZIP-1 plays an important role in defense against intracellular infection, it almost certainly is not the only transcription factor with such a role. Our qRT-PCR and RNA-seq analyses demonstrated that many genes induced as part of the IPR do not require ZIP-1 for their induction, while some require ZIP-1 only for early induction, but not late induction. Future studies with screens for transcription factors that mediate induction of *zip-1*-independent genes should enable a more complete assessment of the immune response to intracellular infection in *C. elegans*.

In this study we demonstrate that ZIP-1 protein expression can be activated by different signaling pathways that act in parallel to induce the IPR, including proteasome inhibition and DRH-1 pathway triggered by viral infection (10). While *zip-1* itself appears to be transcriptionally and translationally induced by infection, we believe that ZIP-1 is the immediate transcription factor that activates IPR gene expression upon various triggers. *zip-1* is required for IPR gene induction only 30 min after activation by bortezomib, which is likely too short a time for a separate transcription factor to activate *zip-1* transcription and translation, which would then induce IPR gene expression. There is still much to be learned about how various triggers activate the IPR, although a likely ligand and receptor pair have been identified for the Orsay virus, where viral RNA replication products appear to be detected by the RIG-I-like receptor DRH-1 (10). As mentioned earlier, *C. elegans* lacks the downstream factors that mediate viral/RIG-I signaling in mammals, such as IRF3/7 and interferon. Therefore, we propose that ZIP-1 and the IPR may play an analogous role to IRF3/7 and interferon in *C. elegans* defense against intracellular infection in intestinal epithelial cells. Further analysis should shed light on how the evolutionarily ancient RIG-I-like receptor family is rewired in *C. elegans* to enable activation of ZIP-1 and downstream defense against intracellular infection.

## Materials and methods

### Worm maintenance

Worms were grown on Nematode Growth Media (NGM) plates seeded with Streptomycin-resistant *E. coli* OP50-1 bacteria at 20°C, unless stated otherwise. Strains used in this study are listed in the Table S5.

### RNAi screens

RNAi screens were performed using the feeding method in liquid medium. Gravid adults were bleached following a standard protocol (65), and isolated eggs were incubated in M9 medium overnight to hatch into starved L1’s unless stated otherwise. In particular, for the screen in the *pals-22(jy3)* mutant background, eggs isolated from bleached gravid adults were put on OP50 seeded NGM plates and incubated at 20°C for 48 h. Subsequently, animals were washed off the plates with S-basal medium and 150 animals were transferred into wells of 96-well plates. Overnight cultures of RNAi HT115 bacterial strains were supplemented with 5 mM isopropyl β-d-1-thiogalactopyranoside (IPTG) and 1 mM carbenicillin, and added to the wells with worms. Control RNAi experiments were carried out using L4440 (negative control vector) and *gfp(RNAi)* (positive control). Following incubation at 20°C for 48 h, animals were collected and analyzed on the COPAS Biosort machine (Union Biometrica). PALS-5::GFP signal and the time-of-flight (TOF, as a measure of length) were quantified, and average values for fluorescence/body length were calculated for each animal. In Table S1 we also list normalized TOF values as a proxy for body length, which indicates that an RNAi clone like *lin-26* had low GFP/TOF, but also had low TOF, indicating small size and thus potentially poor overall health, and thus was not pursued as a hit. Two experimental replicates were performed for majority of RNAi clones (the transcription factor RNAi library is split among five 96-well plates, three of which were tested twice and two of which were tested once).

For the screen in which chronic heat stress was used to induce the IPR, synchronized populations of 150 L1 animals carrying the *jyIs8[pals-5p::gfp]* transgene were transferred to S-basal medium in 96-well plates. The wells were supplemented with overnight RNAi bacterial cultures, as previously described for RNAi screen in *pals-22(jy3)* mutant background. Animals were incubated in the shaker at 20°C for 48 h, and then subjected to chronic heat stress at 30°C for 18 h. Subsequently, *pals-5*p::GFP expression was measured and standardized to the worm length using TOF measurements on the COPAS Biosort machine. Three independent experimental replicates were performed.

### RNAi assays on plates

RNA interference assays were performed using the feeding method. Overnight cultures of HT115 *E. coli* were plated on RNAi plates (NGM plates supplemented with 5 mM IPTG and 1 mM carbenicillin) and incubated at room temperature for 3 days. Synchronized L1 animals were transferred to these plates and incubated at 20°C. Following 48 h incubation, specific phenotypes of animals were analyzed (*pals-5*p::GFP expression after exposure to *zip-1* RNAi; developmental defects after *act-5* RNAi treatment). Control RNAi experiments were carried out using a vector plasmid L4440.

### CRISPR/Cas9-mediated deletions of *zip-1* and *pals-5*

Deletions of *zip-1* and *pals-5* were carried out using the co-CRISPR method with preassembled ribonucleoproteins (66, 67). Cas9-NLS protein (27 µM final concentration) was ordered from QB3 Berkeley; sgRNA components and DNA primers were obtained from Integrated DNA Technologies (IDT).

The following crRNA sequences were used to target *zip-1* gene: acacaggcatctggggaccc (for generating the *jy13* allele), tcagcttgtgctgggcgttg (for generating the *jy14* allele), agcaatttgagccaagctga (for generating both *jy13* and *jy14* alleles). PCR screenings were performed using the primers 1-4 listed in Table S6. Deletion-positive lines were backcrossed three times to the N2 strain before they were used in experiments. The *jy13* allele is an 8241 base pair long deletion, starting 172 nucleotides upstream of the *zip-1* start codon and ending at the last nucleotide before the stop codon (C8069). *jy14* allele is a 4108 base pair long deletion, starting at nucleotide G3962 and ending at the last nucleotide before the stop codon (C8069).

The following crRNA sequences were used to target the *pals-5* gene: aaatactcgaagcaattcag and aaaacgaatagaaaatggga. PCR screenings were performed using primers 10 and 11 from the Table S6. Deletion-positive lines were backcrossed three times to the N2 strain before they were used in experiments. *jy133* allele is a 1706 base pair long deletion, starting 128 nucleotides upstream of the *pals-5* start codon and ending at the 108th nucleotide after the stop codon.

### Orsay virus infections

Orsay virus isolate was prepared as previously described (11). For *pals-5*p::GFP expression analysis and FISH staining for infection level quantification, L1 animals were exposed to a mixture of OP50-1 bacteria and Orsay virus for 12 h at 20°C, whereas animals used for ZIP-1::GFP analysis were infected with a high dose of virus for 9 h at 20°C. *pals-5*p::GFP reporter expression was analyzed in animals that were anesthetized with 10 mM levamisole. For FISH analysis, animals were collected and fixed in 4% paraformaldehyde for 15 to 45 min depending on the assay. Fixed worms were stained at 46°C overnight using FISH probes conjugated to the red Cal Fluor 610 fluorophore, targeting Orsay virus RNA1. GFP imaging and FISH analysis were performed using Zeiss AxioImager M1 compound microscope. For qRT-PCR analyses, synchronized L4 animals were exposed to a mixture of OP50-1 bacteria and Orsay virus for 24 h at 20°C. RNA isolation and qRT-PCR analysis were performed as described below. ZIP-1::GFP expression was analyzed and imaged on a Zeiss LSM700 confocal microscope run by ZEN2010 software.

### Microsporidia infections

*N. parisii* spores were prepared as previously described (63). Spores were mixed with food and L1 synchronized animals (a dose of 8 million spores per plate was used for ZIP-1::GFP expression analyses, a dose of 0.5 million spores per plate was in all other assays). Animals were incubated at 25°C for 3 h (for sporoplasm counting and ZIP-1::GFP analysis), 24 h (for qRT-PCR analysis of IPR gene expression) or 30 h (for pathogen load analysis). For *pals-5*p::GFP expression analysis, animals were anesthetized with 10 µM levamisole and imaged using Zeiss AxioImager M1 compound microscope. For FISH analysis, animals were collected and fixed in 4% paraformaldehyde for 15 to 45 min depending on the assay. Fixed worms were stained at 46°C for 6 h (for ZIP-1::GFP analysis) or overnight (for pathogen load analyses) using FISH probes conjugated to the red Cal Fluor 610 fluorophore, targeting ribosomal RNA. 3 hpi samples were analyzed using Zeiss AxioImager M1 compound microscope; 30 hpi samples were imaged using ImageXpress automated imaging system Nano imager (Molecular Devices, LLC), and fluorescence levels were analyzed using FIJI program. ZIP-1::GFP expression was analyzed and imaged on a Zeiss LSM700 confocal microscope run by ZEN2010 software.

### Bortezomib treatments

Proteasome inhibition was performed using bortezomib (Selleckchem, catalog number S1013) as previously described (18, 22). Synchronized L1 animals were plated on 10 cm (for RNA extraction) or 6 cm NGM plates (for phenotypic analyses and transgene expression measurements), and grown for 44 h or 48 h at 20°C depending on the assay. 10 mM stock solution of bortezomib in DMSO was added to reach a final concentration of 20 µM per plate. The same volume of DMSO was added to the control plates. Plates were dried and worms incubated for 30 minutes, 4, 21 or 25 hours at 20°C. Imaging was performed using Zeiss AxioImager M1 compound microscope or ImageXpress automated imaging system Nano imager (Molecular Devices, LLC), and analyzed using FIJI program. For RNA extraction, animals were washed off the plates using M9, washed with M9 and collected in TRI reagent (Molecular Research Center, Inc.). ZIP-1::GFP expression was analyzed and imaged on a Zeiss LSM700 confocal microscope run by ZEN2010 software.

### Fluorescence measurements

Fluorescence measurements shown in Fig. 1A and B and Fig. 2D were performed using the COPAS Biosort machine (Union Biometrica). The fluorescent signal was normalized to TOF, as a proxy for worm length. Fluorescence measurements shown in Fig. 2F, Fig. 6D and Fig. S12 were performed by imaging animals using ImageXpress automated imaging system Nano imager (Molecular Devices, LLC), followed by image analysis in FIJI. Mean gray value (as a ratio of integrated density and analyzed area) was measured for each animal and normalized to the background fluorescence.

### Bioluminescence measurements

Synchronized L1 animals were grown at 20°C for 44 h and then treated with bortezomib or DMSO for 4 h. Sample preparation and nanoluciferase bioluminescence measurements were performed as previously described (22). In brief, animals were collected and disrupted using silicon carbide beads in lysis buffer (50 mM HEPES pH 7.4, 1 mM EGTA, 1 mM MgCl2, 100 mM KCl, 10% glycerol, 0.05% NP40, 0.5 mM DTT, protease inhibitor cOmplete (Sigma, catalog number 11836170001)). The lysates were centrifuged and the supernatants were collected and stored at -80°C until bioluminescence was measured. Nano-Glo Luciferase Assay System reagent (Promega, catalog number N1110) was added to the worm lysate supernatant before analysis, and incubated at room temperature for 10 minutes. Analysis was performed on a NOVOstar plate reader. The results were normalized to blank controls.

### smFISH analysis

smFISH experiments were performed as previously described (12). In brief, L4 animals were treated with bortezomib or DMSO for 4 h at 20°C. Animals were washed off the plates, fixed in 4% paraformaldehyde in phosphate-buffered saline + 0.1% Tween 20 (PBST) at room temperature for 30 min, and incubated in 70% ethanol overnight at 4°C. Staining was performed with 1 µM Cal Fluor 610 conjugated *pals-5* smFISH probes (Biosearch Technologies) in smFISH hybridization buffer (10% formamide, 2X SSC, 10% dextran sulfate, 2 mM vanadyl ribonucleoside complex, 0.02% RNase free BSA, 50 μg *E. coli* tRNA) at 30°C in the dark overnight. Samples were incubated in the wash buffer (10% formamide, 2X SSC) at 30°C in the dark for 30 min. Vectashield + DAPI was added to each sample, and stained worms were transferred to microscope slides and covered with glass coverslips. Z-stacks of the body region containing anterior part of the intestine was performed using Zeiss AxioImager M1 compound microscope with a 63X oil immersion objective. Image processing was performed using FIJI. smFISH spot quantification was performed using StarSearch program (http://rajlab.seas.upenn.edu/StarSearch/launch.html). When selecting the region of interest, the anterior boundary of the first four intestinal cells was determined based on the prominent border between pharynx and intestine, which is visible in the DIC channel. The posterior boundary was set at the middle distance between DAPI-stained nuclei of the first and the second intestinal rings.

### PALS-5 expression and anti-PALS-5 antibody synthesis

A *pals-5* cDNA with N-terminal sequence (5’-tatgcatcaccaccatcaccatgaaaatctgtattttcag-3’) and C-terminal sequence (5’-gagagaccggccggccgatccggctgctaa-3’) was synthesized as a gBlock (Genewiz) and cloned into BsaI-HFv2 digested into a custom vector derived from pET21a. The resulting plasmid (pBEL2159), which includes an N-terminal His-TEV-tag, was transformed into Rosetta (DE3) cells (Novagen) for protein expression. For expression, LB with carbenicillin/ chloramphenicol was inoculated with Rosetta (DE3)/pBEL2159 and grown at 37°C with shaking at 200rpm. The overnight culture was diluted 1:50 in LB+carbenicillin/chloramphenicol and then induced by adding IPTG to a final concentration of 1mM at 16°C, and allowed to shake overnight. Cells were harvested by centrifugation and resuspended in lysis buffer (50mM Tris pH8, 300mM NaCl, 10mM Imidazole, 10% Glycerol, 1mM phenylmethylsulfonyl fluoride (PMSF)). Cells were lysed using the Emulsiflex-C3 cell disruptor (Avestin) and then centrifuged at 4°C, 12,000g to pellet cell debris. The pellet, containing a large amount of insoluble PALS-5, was resuspended in urea lysis buffer (100 mM NaH2PO4/10 mM Tris base, 10 mM Imidazole, 8 M Urea [titrated to pH8 by NaOH]). The solubilized pellet was centrifuged at 4000g, and the supernatant collected. PALS-5 from the resulting supernatant was passed twice through NiNTA resin (Cytiva #17531802), which was subsequently washed with urea wash buffer (100 mM NaH2PO4/10 mM Tris base, 40 mM Imidazole, 8 M Urea [titrated to pH8 with NaOH]), and the bound proteins were then eluted in urea elution buffer (100 mM NaH2PO4/10 mM Tris base, 300 mM Imidazole, 8 M Urea [titrated to pH8 by NaOH]). Fractions containing PALS-5 were pooled, concentrated and dialyzed into dialysis buffer (PBS (137mM NaCl, 2.7mM KCl, 1.5mM KH2PO4, 8.1mM Na2HPO4), 3.9M Urea [titrated to pH8 by NaOH]) overnight at room temperature. The following day, the sample was dialyzed once again in fresh dialysis buffer for 3hr at room temperature. The dialyzed sample was supplemented with 10% glycerol, flash frozen in liquid nitrogen for storage, and submitted to ProSci Inc. for custom antibody production (Poway, CA). Rabbits were initially immunized with 200 µg full-length His::TEV tagged PALS-5 antigen in Freund’s Complete Adjuvant. Rabbits were then subsequently boosted with four separate immunizations of 100 µg antigen in Freund’s Incomplete Adjuvant over a 16-week period. Approximately 25 ml of serum was collected and PALS-5 polyclonal antibody was purified with an immuno-affinity chromatography column by cross-linking PALS-5 to cyanogen bromide (CNBr)-activated Sepharose 4B gel. Antibody was eluted from the affinity column in 100 mM glycine buffer pH 2.5, precipitated with polyethylene glycol (PEG), and concentrated in PBS pH 7.4 + 0.02% sodium azide. Antibody concentration was determined by ELISA and used in western blot analysis described below.

### Western blot analysis

3000 L4 animals were treated with bortezomib or DMSO for 4 h at 20°C, and then collected and washed with M9. 20 µl of 6x loading buffer (375 mM Tris⋅HCl pH 6.8, 600 mM DTT, 12% SDS, 0.06% bromophenol blue, and 60% glycerol) were added to the final sample volume of 100 µl. Samples were boiled at 100°C for 10 min and stored at -30°C. Proteins were separated on a 10% sodium dodecyl sulfate–polyacrylamide gel electrophoresis precast gel (Bio-Rad), and transferred onto polyvinylidene difluoride (PVDF) membrane. 5% nonfat dry milk in PBST was used to block for nonspecific binding for 2 h at room temperature. The membranes were incubated with primary antibodies overnight at 4°C (rabbit anti-PALS-5 diluted 1:1,000 and mouse anti-tubulin (Sigma, catalog number T9026) diluted 1:3000 in blocking buffer). Next, the membranes were washed five times in PBST, and then incubated in horseradish peroxidase-conjugated secondary antibodies at room temperature for 2 h (goat anti-rabbit (MilliporeSigma, catalog number 401315) and goat anti-mouse (MilliporeSigma, catalog number 401215) diluted 1:10,000 in blocking buffer). After five washes in PBST, the membranes were treated with enhanced chemiluminescence (ECL) reagent (Amersham) for 5 min, and imaged using a Chemidoc XRS+ with Image Lab software (Bio-Rad). Quantification of band intensities in 3 Western blot replicates was performed using Image Lab software (Bio-Rad). PALS-5 band intensities for each sample were normalized to the ratio of the tubulin expression levels between N2 DMSO control and a given sample.

### RNA isolation

Total RNA isolation was performed as previously described (15). Animals were washed off plates using M9, then washed with M9 and collected in TRI reagent (Molecular Research Center, Inc.). RNA was isolated using BCP phase separation reagent, followed by isopropanol and ethanol washes. For RNA seq analysis, samples were additionally purified using RNeasy Mini kit from Qiagen.

### qRT-PCR analyses

qRT-PCR analysis was performed as previously described (15). In brief, cDNA was synthesized from total RNA using iScript cDNA synthesis kit (Bio-Rad). qPCR was performed using iQ SYBR Green Supermix (Bio-Rad) with the CFX Connect Real-Time PCR Detection System (Bio-Rad). At least three independent experimental replicates were performed for each qRT-PCR analysis. Each sample was analyzed in technical duplicates. All values were normalized to expression of *snb-1* control gene, which does not change expression upon IPR activation. The Pffafl method was used for data quantification (68). The sequences of the primers used in all qRT-PCR experiments are given in the Table S6 (primers 12-33).

### RNA seq analysis

cDNA library preparation and single-end sequencing was performed at the Institute for Genomic Medicine at the University of California, San Diego. Reads were mapped to *C. elegans* WS235 genome using Rsubread in RStudio (Tables S7 and S8). Differential expression analyses were performed using limma-voom function in Galaxy platform (https://usegalaxy.org/). Genes with counts number lower than 0.5 counts per million (CPM) for 30 min timepoint samples and 1 CPM for 4 h timepoint samples were filtered out. Quality weights were applied in analysis of 30 min timepoint. Differentially expressed genes had adjusted *p*-value lower than 0.05. Visualization of the mapped reads shown in the Fig. S9 was performed using Integrative Genomics Viewer (Broad Institute) (69).

### Analysis of enriched gene categories in *zip-1*-dependent gene datasets

Annotation and visualization of genes upregulated in wild-type but not in *zip-1(jy13)* background was performed using WormCat online tool (http://www.wormcat.com/) (26).

### Comparisons of differentially expressed genes from different datasets

An R studio package GeneOverlap was used for RNA-seq datasets comparative analyses. Differentially expressed genes from RNA-seq analyses from this study were compared with relevant previously published datasets (11, 13, 15, 24, 25, 34, 70–72). Statistical similarity between datasets was determined using Fisher’s exact test. The odds ratios, Jaccard indexes and *p*-values were calculated. Total number of genes was set to 46902. Data are represented in the contingency tables in which odds ratio and Jaccard index values are shown in the heat map format, whereas *p*-values are indicated numerically.

### CRISPR/Cas9-mediated tagging of *zip-1*

A long, partially single-stranded DNA donor CRISPR-Cas9 method was employed to endogenously tag the *zip-1* locus (73). A single sgRNA (agcaatttgagccaagctga) was used to preassemble ribonucleoprotein with Cas9 (IDT). Repair templates that contain *gfp*, *sbp* (Streptavidin-Binding Peptide) and *3xFlag* tags were amplified from plasmid pET386 using primers 5-8 from Table S6. Injection quality was monitored by co-injecting animals with pRF4 plasmid (*rol-6(su1006)* marker). PCR screening of GFP insertion was performed using primers 3, 4 and 9 from the Table S6. A line containing *gfp::sbp::3xFlag* insertion before endogenous *zip-1* stop codon was backcrossed three times to the N2 strain before it was used in experiments.

### Tissue-specific RNAi analysis

Tissue-specific RNAi analysis was performed using the feeding method. *E. coli* OP50-1 strain was modified to enable *zip-1* RNAi or control RNAi (L4440). Bacterial overnight cultures were plated on NGM plates supplemented with 2.2 mM IPTG and 1 mM carbenicillin, and incubated at room temperature for 3 or 4 days. 3000 synchronized L1 animals were transferred to prepared plates and grown at 20°C for 48 h. Animals were then treated with bortezomib or DMSO as described earlier. VP303 (*rde-1*) and MGH167 (*sid-1*) strains were used for intestinal RNAi; NR222 (*rde-1*) strain was used for epidermal RNAi. Replicates that were included into analysis of *sid-1* mutants had at least 50-fold increase in *pals-5* expression levels on control RNAi plates following bortezomib treatment. This threshold allowed detection of any substantial decrease in *pals-5* induction in *zip-1(RNAi)* samples.

### Killing assays

For *N. parisii* killing assays, about 150 L1 worms were mixed with 50 μl of a 10X concentration of OP50-1 *E. coli* and 1 million *N. parisii* spores, and placed onto a 3.5 cm tissue culture-treated NGM plate (3 plates for each strain). After 66 h of infection at 25°C, alive animals were transferred onto new NGM plates containing only OP50-1 *E. coli* food (30 animals per plate, 3 plates per worm strain). Animals were scored daily and alive animals were transferred to fresh NGM plates. Data from 3 experimental replicates were merged and analyzed using Survival function in GraphPad Prism 9; log-rank (Mantel-Cox) test was used for statistical analyses.

### Longevity assays

For longevity assays, about 75 L1 worms were mixed with 50 μl of a 10X concentration of OP50-1 *E. coli*, and placed onto a 3.5 cm tissue culture-treated NGM plate (3 plates for each strain). After 66 h incubation at 25°C, animals were transferred to new NGM plates supplemented with OP50-1 *E. coli* food source (30 animals per plate, 3 plates per strain). Animals were scored daily and alive animals were transferred to fresh NGM plates. Data from 3 experimental replicates were merged and analyzed using Survival function in GraphPad Prism 9; log-rank (Mantel-Cox) test was used for statistical analyses.

### Bead feeding assay

2000 synchronized L1 worms were mixed with 6 μl fluorescent beads (Fluoresbrite Polychromatic Red Microspheres, Polysciences Inc.), 25 μl 10X concentrated OP50 *E. coli*, 500.000 *N. parisii* spores and M9 (total volume 300 ul). This mixture was then plated on 6 cm NGM plates, allowed to dry for 5 min and then incubated at 25°C. After 5 min, plates were shifted to ice, washed with ice-cold PBST and fixed in 4% paraformaldehyde. Animals were imaged using ImageXpress automated imaging system Nano imager (Molecular Devices, LLC). Fluorescence was analyzed in FIJI program.

### Thermotolerance assay

Animals were grown on NGM plates at 20°C until L4 stage. L4 animals were transferred to new plates and exposed to heat shock at 37.5°C for 2 h. Recovery was performed at room temperature for 1 h on a single layer, followed by 24 h incubation at 20°C. After this time, animals were scored for viability based on their ability to move after touch. Three plates with 30 animals per plate were analyzed for each strain. Three experimental replicates were performed.

### Body length measurements

For body length analysis of wild-type and *sid-1*(*-*)*; vha-6p::sid-1* mutant strains (Fig. S10B), synchronized L1 animals were placed on control or *act-5* RNAi plates and allowed to grow at 20°C for 48 h. For analysis of wild type, *zip-1(jy13)* and *pnp-1*(*-*) mutants (Fig. S13B), synchronized L1 animals were plated on NGM plates and allowed to grow at 20°C for 44 h. Animals were washed off the plates with M9 and fixed in 4% paraformaldehyde (Fig. S10B) or anesthetized with 10 µM levamisole (Fig. S13B). Animals were imaged using ImageXpress automated imaging system Nano imager (Molecular Devices, LLC) in 96-well plates. Length of each animal was measured using FIJI program. 50 animals were analyzed for each strain, in each of three experimental replicates.

### Data availability

RNA-seq reads were uploaded to the NCBI GEO database with Accession number GSE183361. All data supporting this manuscript is available from the corresponding author upon request.

## Supplementary table legends

**Table S1. Results of RNAi screens.** The expression of PALS-5::GFP reporter was analyzed in *pals-22(jy3)* mutant background. Expression of *pals-5*p::GFP reporter was analyzed in animals exposed to prolonged heat stress. The values of GFP intensity were normalized to the length of worms (TOF).

**Table S2. An overview of differentially expressed genes in animals treated with bortezomib and DMSO.** Differentially expressed genes with adjusted *p*-value lower than 0.05 are listed for wild-type (N2) animals and *zip-1(jy13)* mutants.

**Table S3. Comparisons of differentially expressed genes from different datasets.** Differentially expressed genes form previously published datasets and their overlap with *zip-1*-dependent genes are shown.

**Table S4. Wormcat analysis results.** Overrepresented categories are listed for both analyzed time points. All catalog values represent number of genes in a specific category in the whole annotation list. Bonferroni values represent corrected *p*-values (Bonferroni correction).

**Table S5. List of worm strains used in this study.** Names of strains and their genotypes are listed.

**Table S6. List of primers used in this study.** Primer labels, descriptions and sequences are listed.

**Table S7. RNA-seq statistics.** Numbers of total and mapped reads are given for each sample and each replicate. R1, R2 and R3 represent replicate 1, 2 and 3 respectively.

**Table S8. Normalized counts for all mapped genes and samples from RNA-seq analysis.**

## Acknowledgements

This work was supported by NIH under R01 AG052622 and GM114139 to ERT, NIGMS/NIH award K12GM068524 to SSG and the American Heart Association postdoctoral award 19POST34460023 to VL. We thank Damian Ekiert, Crystal Chhan, Eillen Tecle, and Cheng-Ju Kuo for helpful comments on the manuscript. We thank Eillen Tecle for crossing strains to create *zip-1(jy13); pnp-1; jyIs8* mutant and for performing preliminary analyses on these animals. We thank Damian Ekiert for his help with PALS-5 protein synthesis. We thank Yishi Jin and Rose Malinow for providing reagents. RNA-seq data were generated at the UC San Diego IGM Genomics Center utilizing an Illumina NovaSeq 6000 that was purchased with funding from a National Institutes of Health SIG grant (#S10 OD026929). The models in Fig. 7 were created using BioRender.com.

## Figures and figure legends

**Fig. S1.**
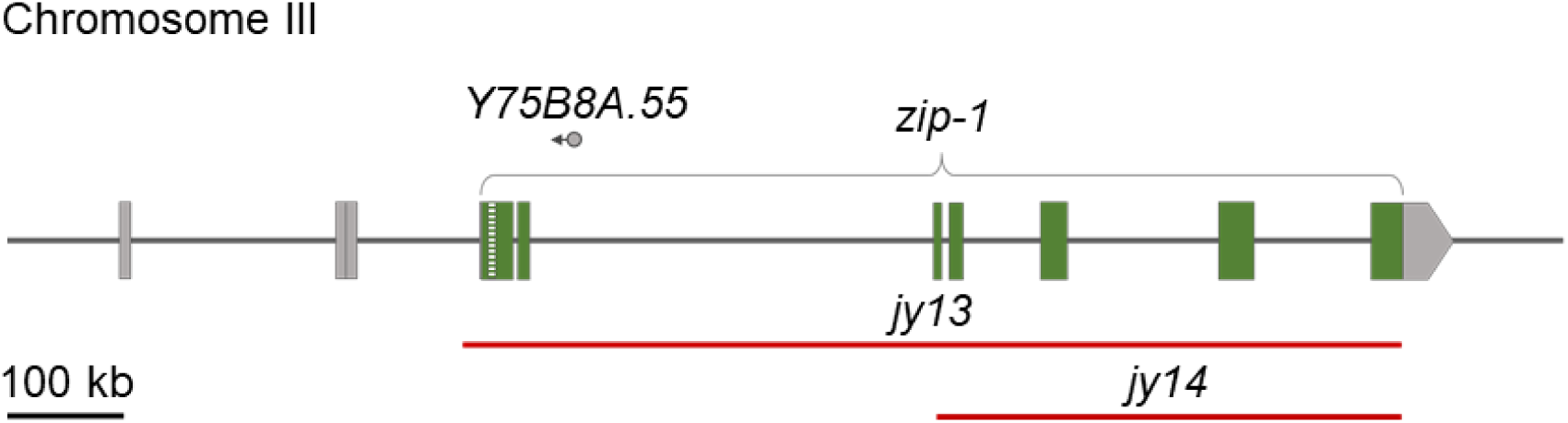
Graphic representation of the *zip-1* gene. Green boxes indicate exons; the box with green stripes represents part of the gene that is spliced in some *zip-1* isoforms. Grey boxes represent 5’ UTR regions annotated for different *zip-1* isoforms, as well as 3’ UTR. Red lines indicate regions deleted in *jy13* and *jy14* alleles. *Y75B8A.55* non-coding RNA is indicated with a circle and an arrow.

**Fig. S2.**
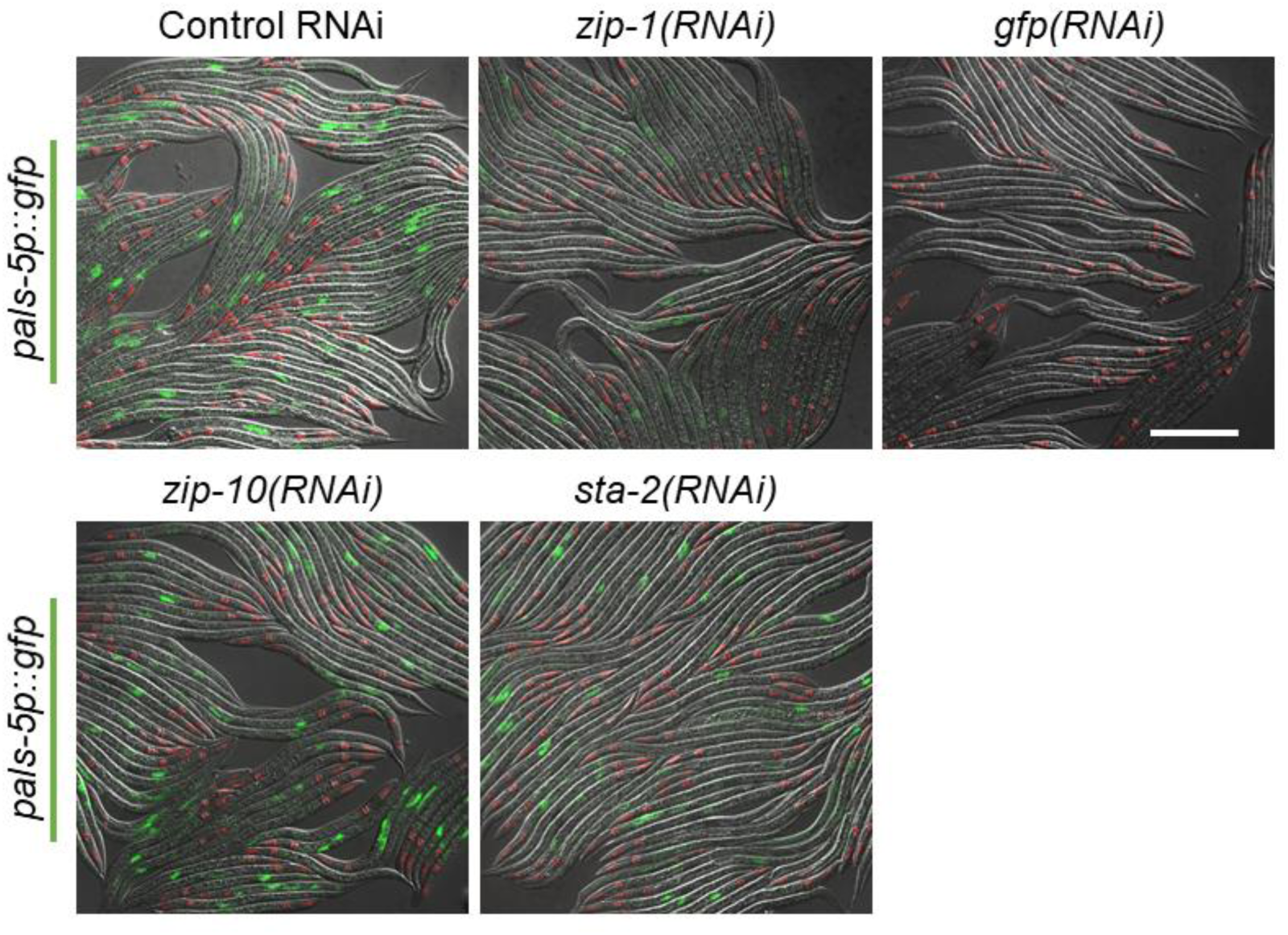
*pals-5*p::GFP expression in animals treated with *zip-10* and *sta-2* RNAi followed by prolonged heat stress. Fluorescent and DIC images were merged. *myo-2*p::mCherry is expressed in the pharynx and is a marker for the presence of the *jyIs8* transgene. Scale bar = 200 µm.

**Fig. S3.**
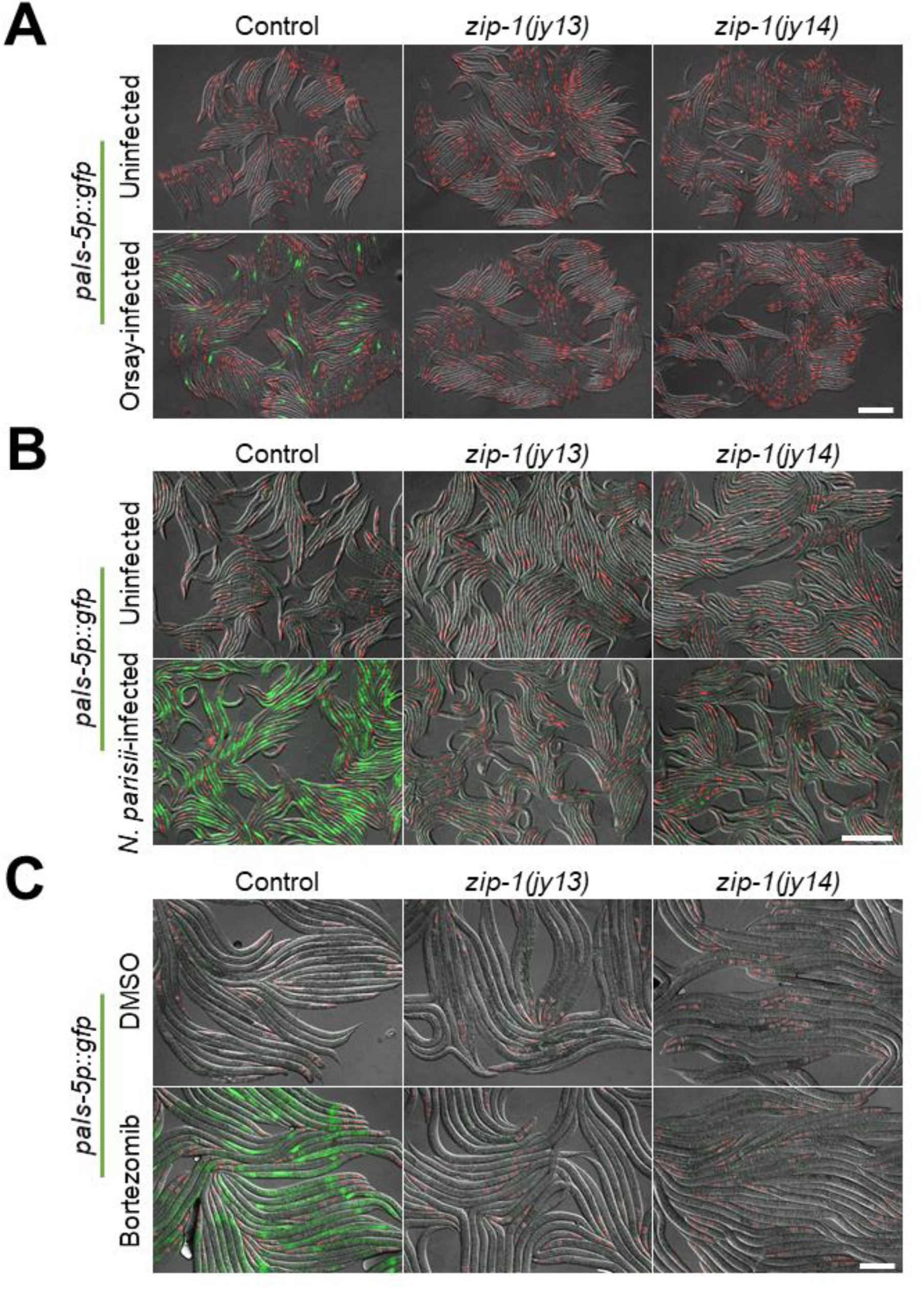
Induction of *pals-5*p::GFP expression is reduced in *zip-1(jy14)* mutants following intracellular infection with Orsay virus or *N. parisii*, as well as following proteasome inhibition by bortezomib. (A-C) Representative images of control (upper row) and Orsay virus-infected (A), *N. parisii*-infected (B) and bortezomib-treated animals (C) (lower row). *myo-2*p::mCherry is expressed in the pharynx and is a marker for the presence of the *jyIs8* transgene. Fluorescent and DIC images were merged. Scale bars = 200 µm.

**Fig. S4.**
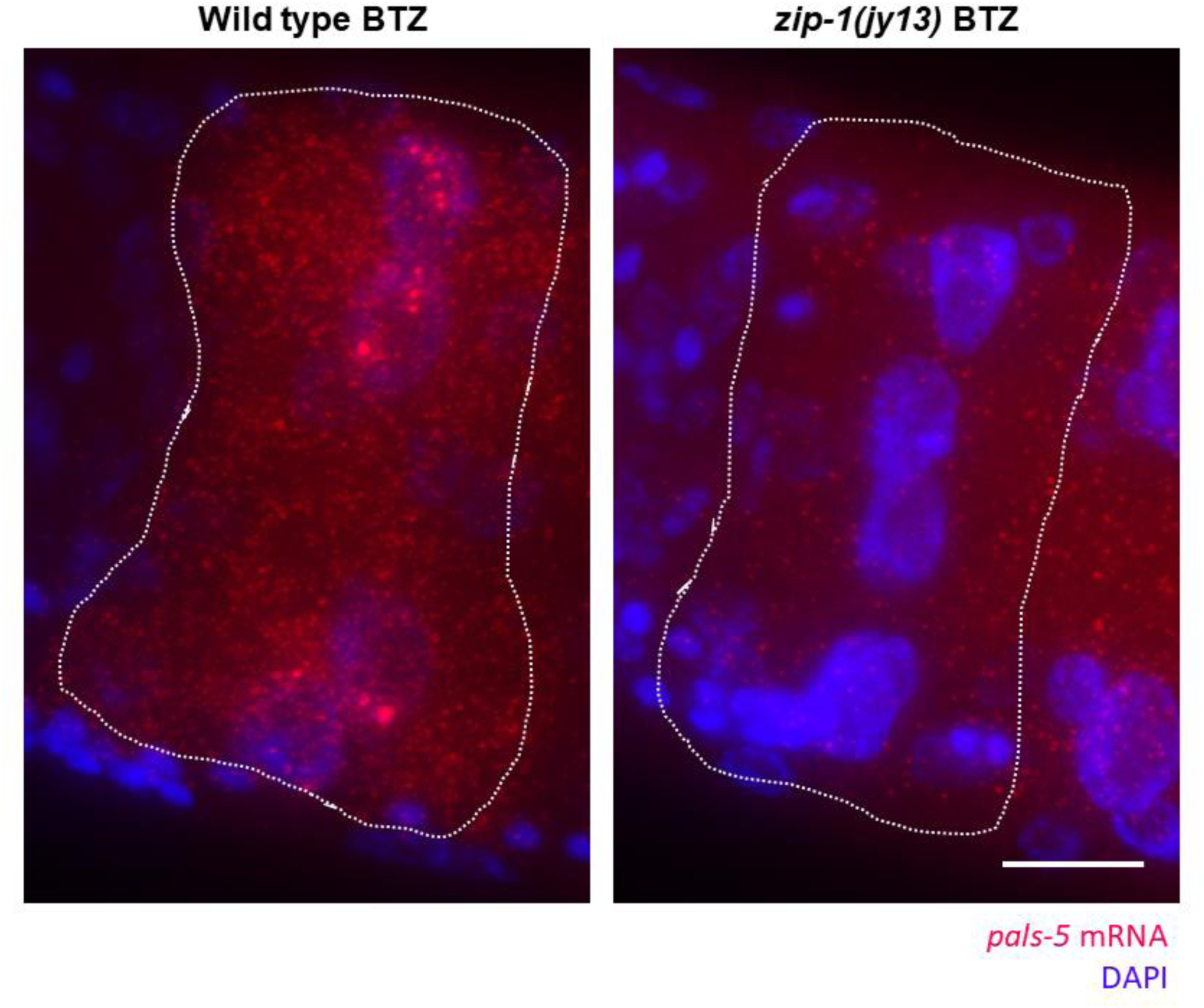
Representative images of the first four intestinal cells of bortezomib treated animals from smFISH analysis. *pals-5* mRNA is visualized with far-red fluorophore and nuclei are labeled with DAPI (blue). Images are maximal projections of z-stacks taken in far-red and blue channels. Dotted lines demarcate areas of the first four intestinal cells that were analyzed. Scale bar = 10 µm.

**Fig. S5.**
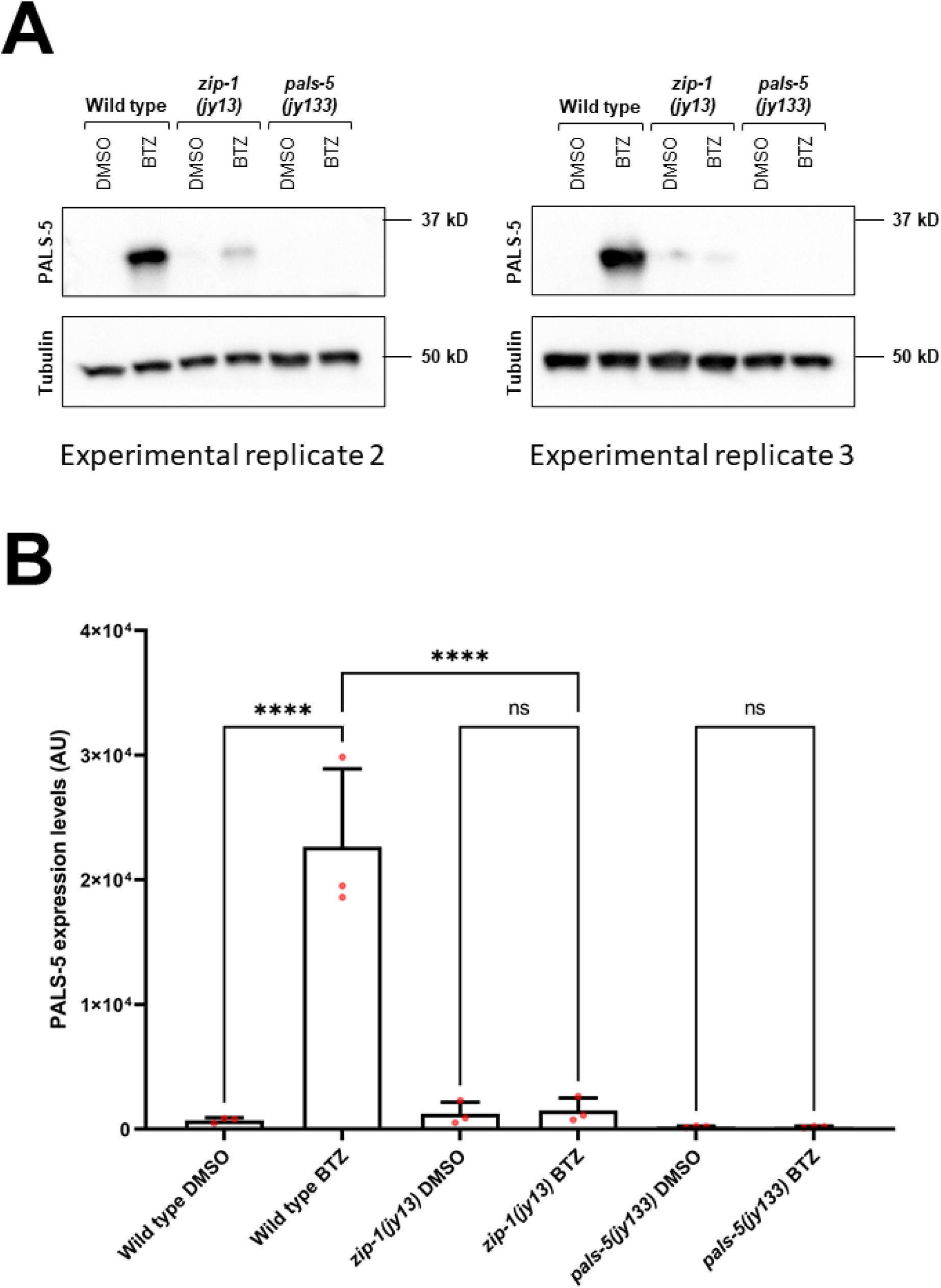
Western blot analysis of PALS-5 expression in wild-type, *zip-1(jy13)* and *pals-5(jy133)* animals. (A) Western blot images of the second and third experimental replicates of PALS-5 protein expression analysis in wild-type, *zip-1(jy13)* and *pals-5(jy133)* animals treated with DMSO and bortezomib. PALS-5 was detected using anti-PALS-5 antibody, whereas anti-tubulin antibody was used as a loading control. Predicted sizes are 35.4 kD for PALS-5 and around 50 kD for different members of tubulin family. (B) Graphical representation of PALS-5 protein expression levels in wild-type, *zip-1(jy13)* and *pals-5(jy133)* animals treated with DMSO and bortezomib. Bar height indicates mean expression value for each sample; red dots indicate values from each of three experimental replicates; error bars represent standard deviations. Strain and treatment information are indicated on x axis; PALS-5 protein levels normalized to tubulin levels are indicated on y axis in arbitrary units (AU). Statistical analysis was performed using an ordinary one-way ANOVA test to calculate *p*-values; **** *p* < 0.0001; ns indicates nonsignificant difference (*p* > 0.05).

**Fig. S6.**
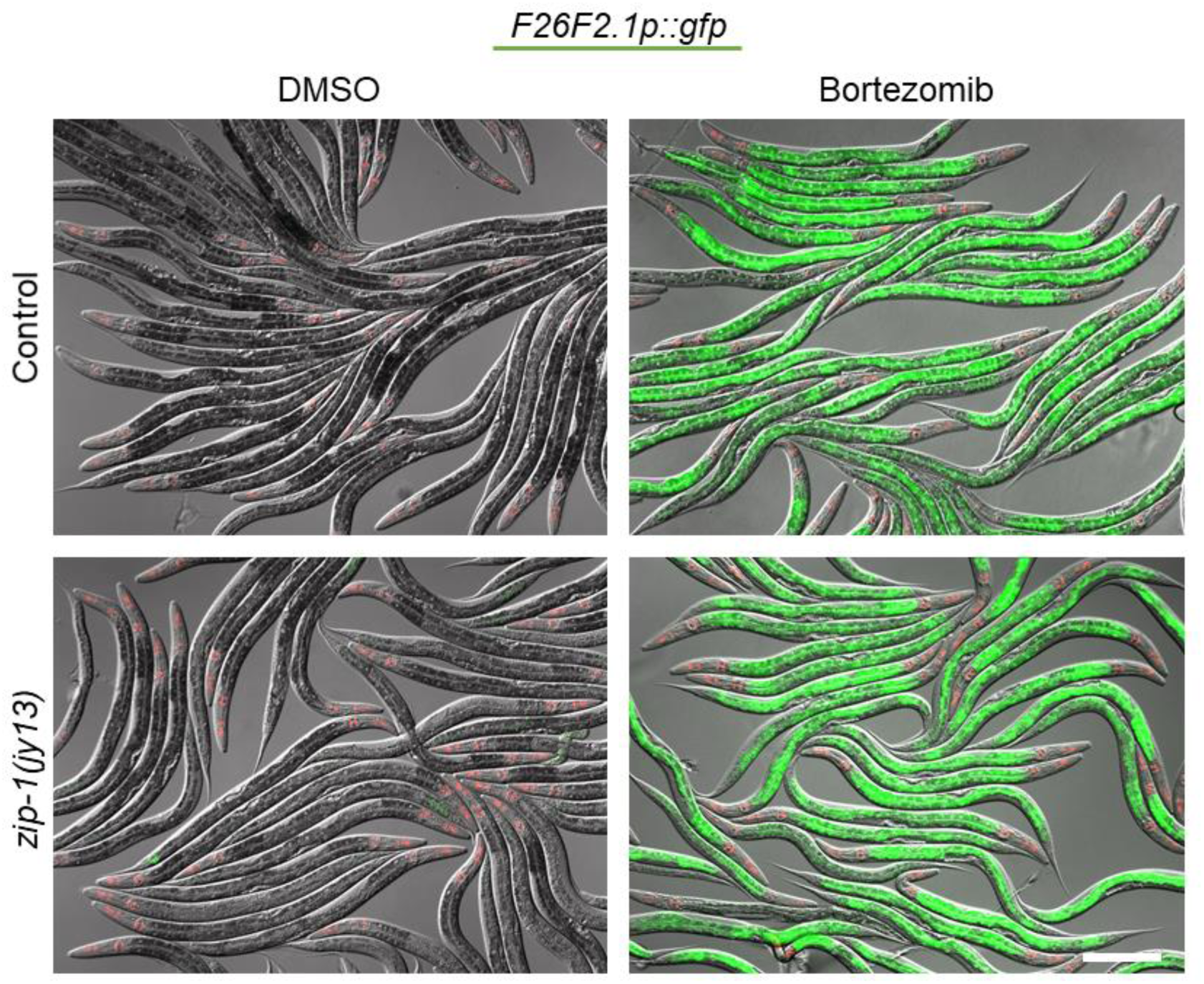
Proteasome inhibition by bortezomib induces *F26F2.1p::*GFP expression in a *zip-1(jy13)* background. Fluorescent and DIC images were merged. *myo-2*p::mCherry is expressed in the pharynx and is a marker for the presence of the *jyIs8* transgene. Scale bar = 200 µm.

**Fig. S7.**
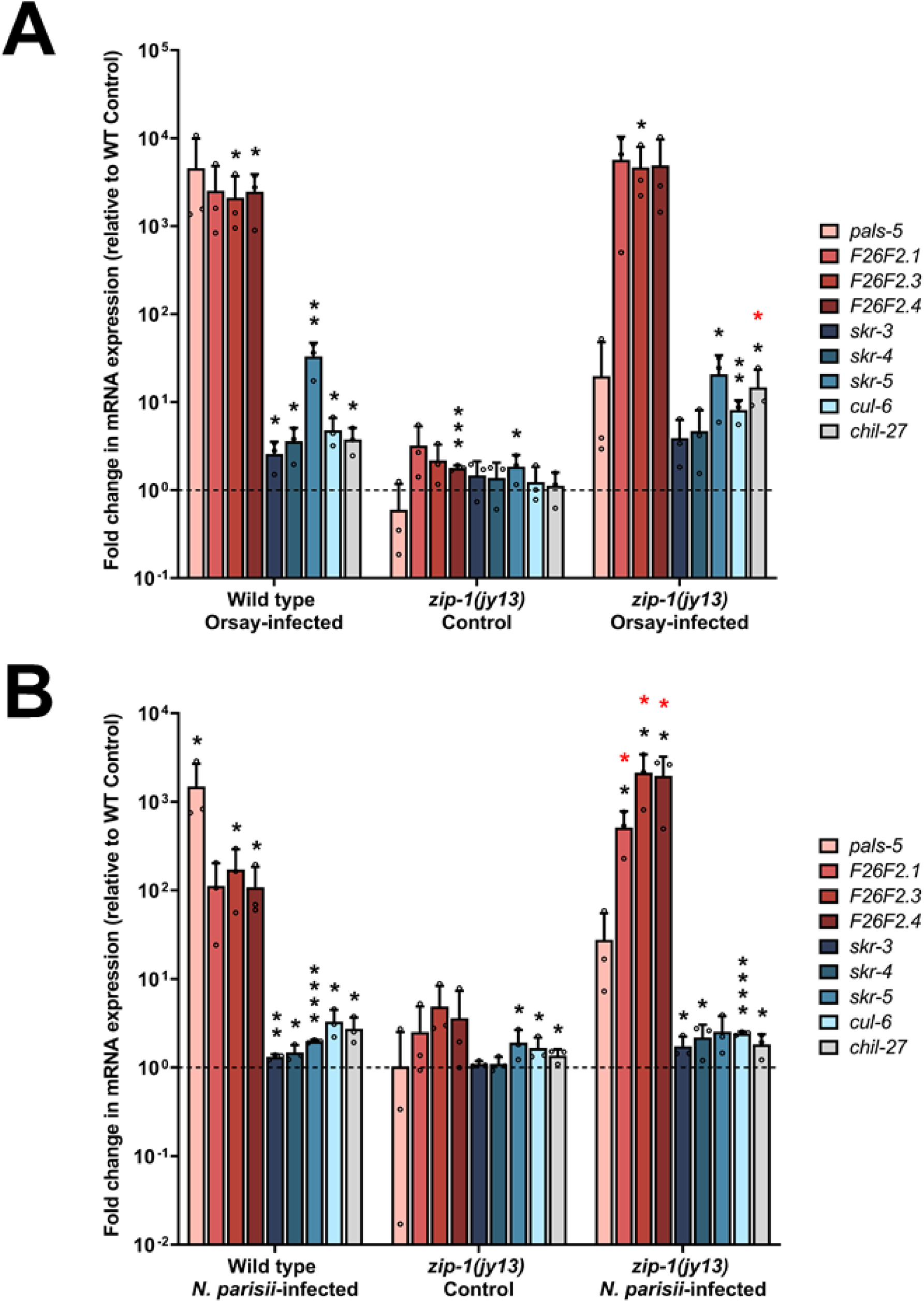
*zip-1* regulates expression of some IPR genes following intracellular infection. (A, B) qRT-PCR measurements of selected IPR genes and *chil-27* in wild-type and *zip-1(jy13)* animals following Orsay virus (A) and *N. parisii* infection (B). The results are shown as the fold change in gene expression relative to uninfected control strain. Three independent experimental replicates were analyzed, the values for each replicate are indicated with circles. Error bars represent standard deviations. A one-tailed t-test was used to calculate *p*-values; black asterisks represent significant difference between the labeled sample and the uninfected wild-type control; red asterisks represent significant difference between infected wild-type and infected *zip-1(jy13)* samples; **** *p* < 0.0001; *** *p* < 0.001; ** *p* < 0.01; * 0.01 < *p* < 0.05; *p*-values higher than 0.05 are not labeled.

**Fig. S8.**
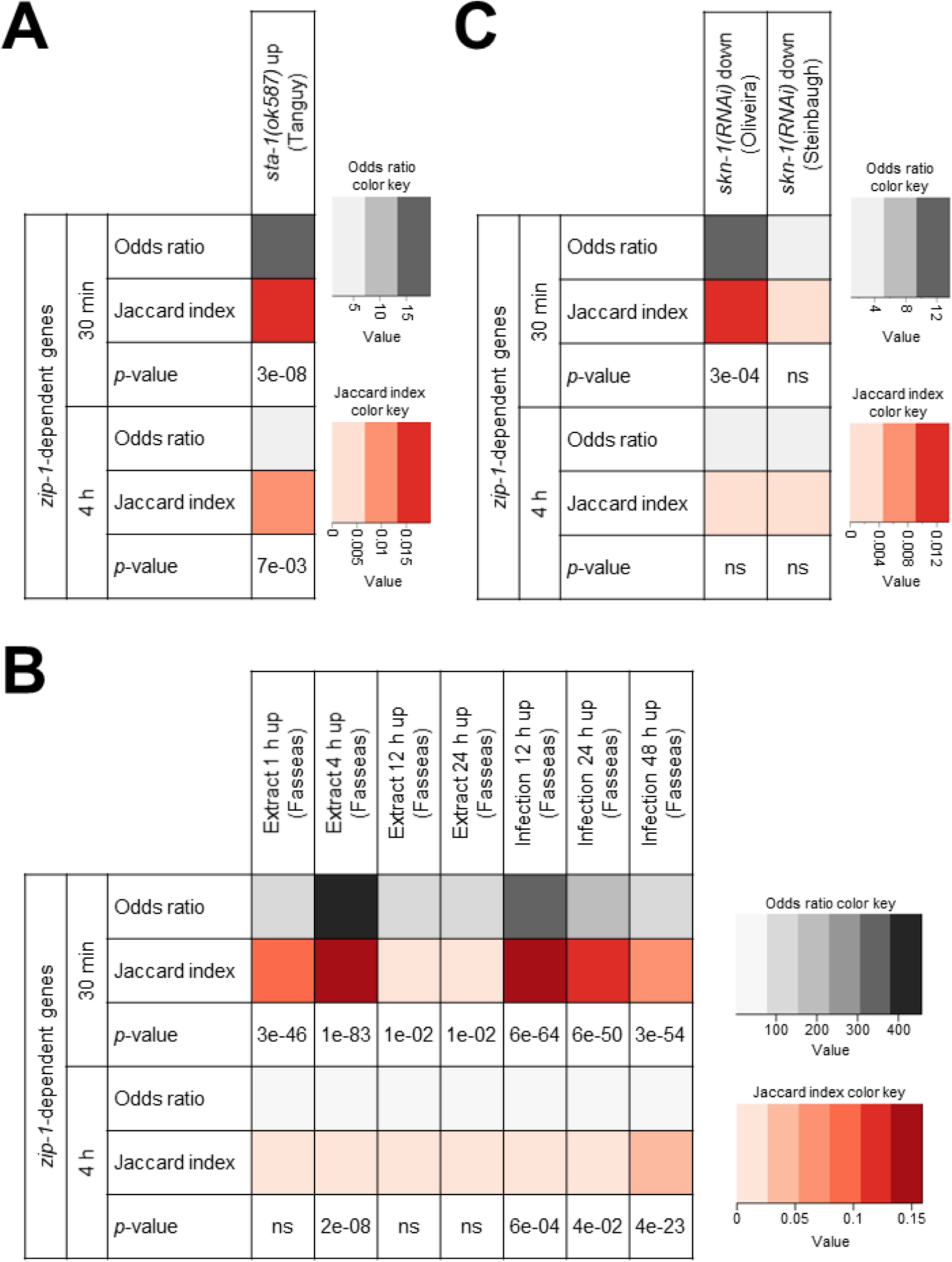
Correlation between *zip-1*-dependent genes and *sta-1*-regulated, ORR and *skn-1*-regulated genes. (A-C) Statistical similarity between *zip-1*-dependent gene set and genes downregulated in *sta-1(ok587)* mutants (A), ORR genes (B) and genes upregulated following *skn-1* downregulation (C). Fisher’s exact test was used to calculate odds ratios and *p*-values. If odds ratio is greater than one, two data sets are positively corelated. Jaccard index measures similarity between two sets, with the range 0-1 (0 – no similarity, 1 – same datasets). For approximate quantification, the odds ratio and Jaccard index color keys are indicated on the right side of each table.

**Fig. S9.**
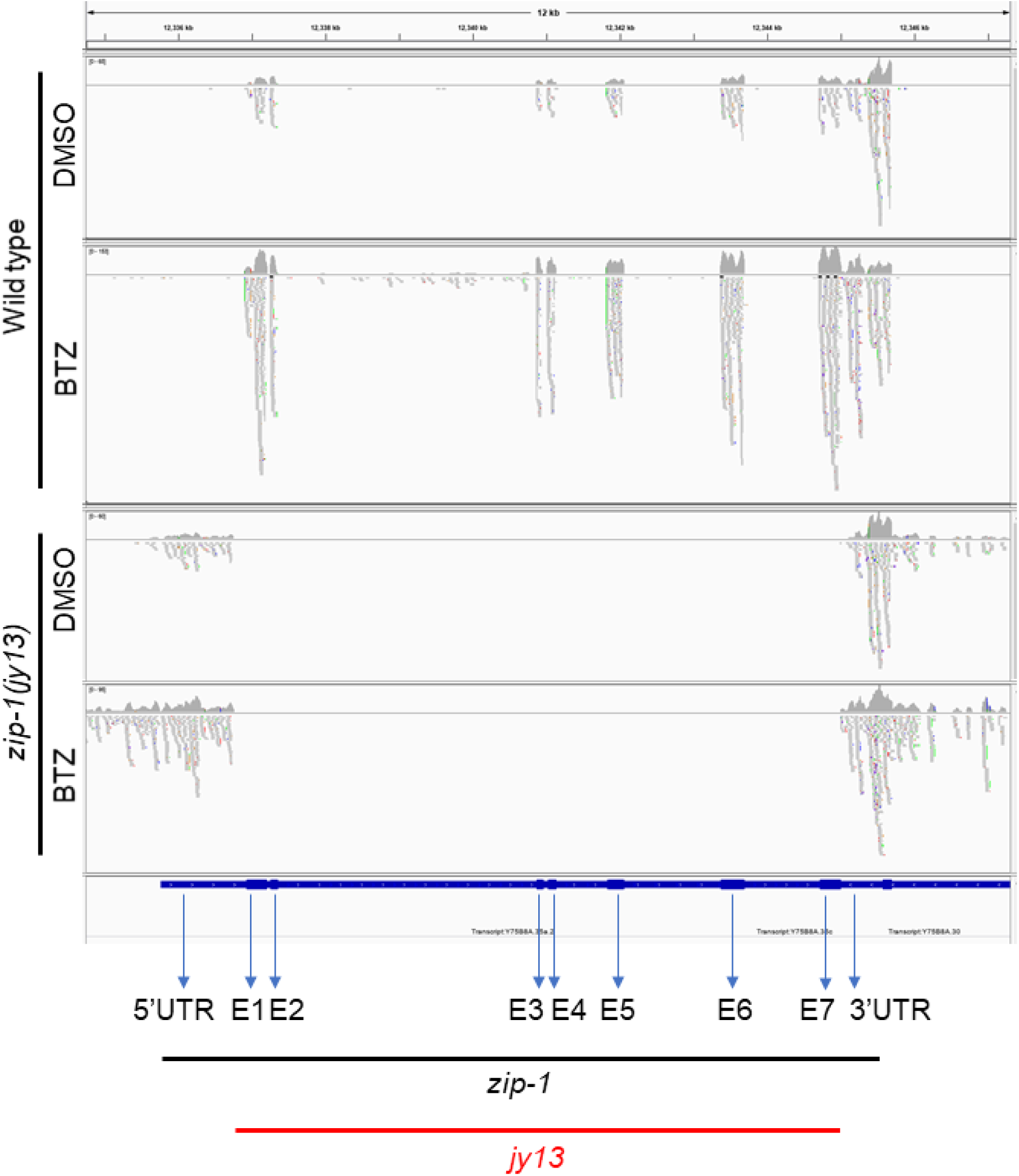
Alignment of mapped *zip-1* reads from RNA seq analysis. Individual mapped reads and summary graphs are shown for DMSO and bortezomib treated wild-type N2 and *zip-1(jy13)* samples. Genomic location is indicated on the top of the graph. *zip-1* and *jy13* locations are indicated on the bottom. Exons of *zip-1* are labeled with E1-E7.

**Fig. S10.**
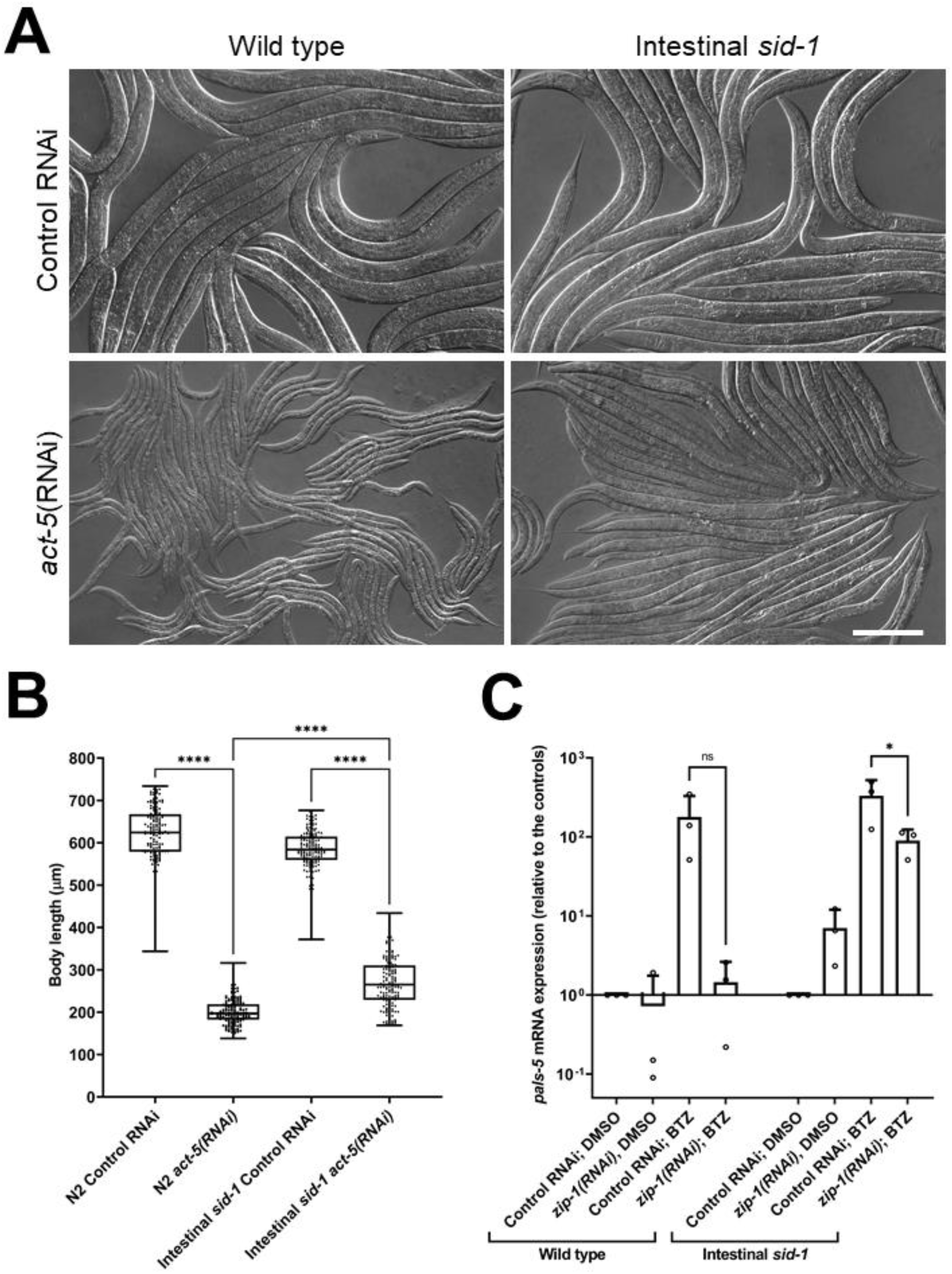
Intestine-specific *zip-1(RNAi*) (*sid-1* background) reduces *pals-5* mRNA induction. (A) Representative DIC images of wild-type and *sid-1* mutant animals following control and *act-5* RNAi treatments. Scale bar = 200 µm. (B) Graphical representation of body length measurements after 48 h incubation at 20°C. 150 animals were analyzed, 50 animals per each of three replicates. The box-and-whisker plots were used for data representation. Each box represents 50% of the data closest to the median value (line in the box). Whiskers span the values outside of the box. (C) qRT-PCR measurements of *pals-5* levels at the 30 min timepoint of bortezomib (BTZ) or DMSO treatments. The results are shown as fold change in gene expression relative to DMSO diluent control. Three independent experimental replicates were analyzed; the values for each replicate are indicated with circles. Error bars represent standard deviations. (B, C) A Kruskal-Wallis (B) and a one-tailed t-test (C) were used to calculate *p*-values; **** *p* < 0.0001; * 0.01 < *p* < 0.05; ns indicates nonsignificant difference (p > 0.05).

**Fig. S11.**
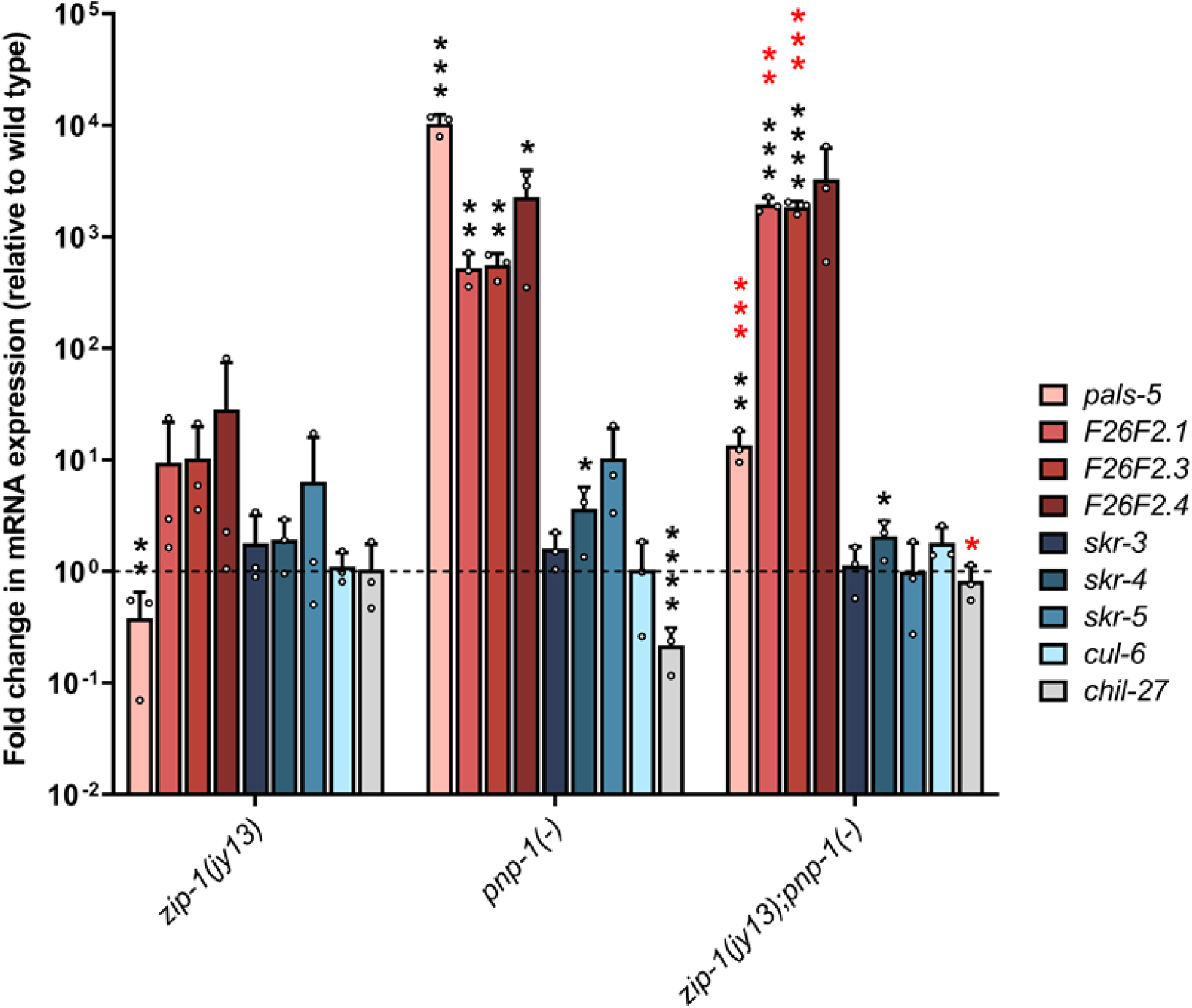
*zip-1* regulates expression of some IPR genes that are upregulated in *pnp-1(jy90)* mutants. qRT-PCR measurements of selected IPR genes and *chil-27* in wild-type, *zip-1(jy13)*, *pnp-1*(*-*) and *zip-1(jy13); pnp-1*(*-*) animals. The results are shown as the fold change in gene expression relative to control strain. All strains are in *jyIs8[pals-5p::gfp; myo-2p::mCherry]* strain background. Three independent experimental replicates were analyzed, the values for each replicate are indicated with circles. Error bars represent standard deviations. A one-tailed t-test was used to calculate *p*-values; black asterisks represent significant difference between the labeled sample and the wild-type control; red asterisks represent significant difference between *pnp-1(jy90)* and *zip-1(jy13); pnp-1(jy90)* backgrounds; **** *p* < 0.0001; *** *p* < 0.001; ** *p* < 0.01; * 0.01 < *p* < 0.05; *p*-values higher than 0.05 are not labeled.

**Fig. S12.**
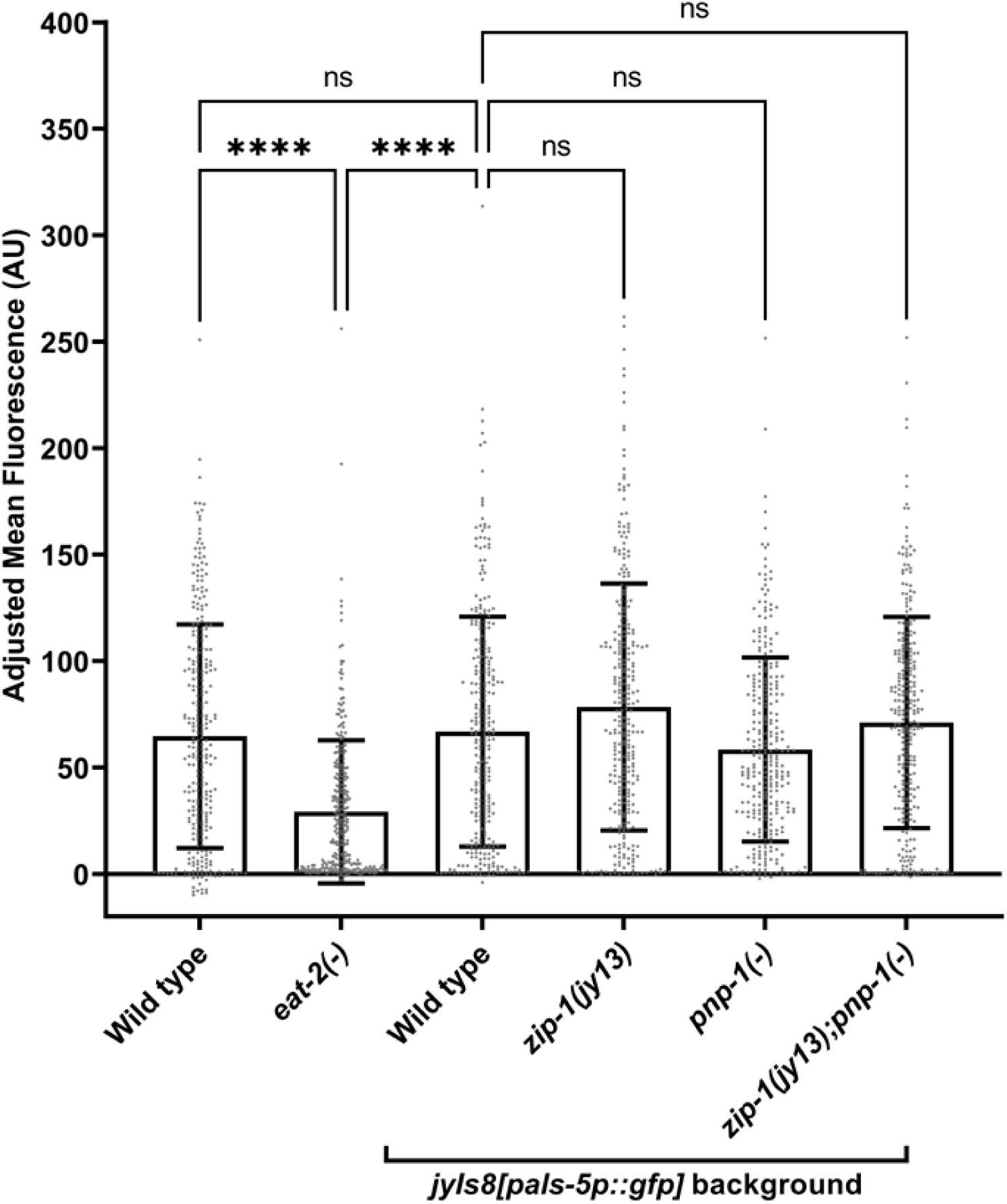
*zip-1(jy13)* and *pnp-1(jy90)* single and double mutants have similar accumulation of fluorescent beads. Quantification of fluorescent bead accumulation in the control strains, *zip-1(jy13); jyIs8*, *pnp-1(jy90); jyIs8* and *zip-1(jy13); pnp-1(jy90); jyIs8* mutants. Mean fluorescence was measured in 150 animals per genotype; background fluorescence was subtracted. In the box-and-whisker plot, each box represents 50% of the data closest to the median value (line in the box). Whiskers span the values outside of the box. AU – arbitrary units. A Kruskal-Wallis test was used to calculate *p*-values; **** *p* < 0.0001; ns indicates nonsignificant difference (*p* > 0.05).

**Fig. S13.**
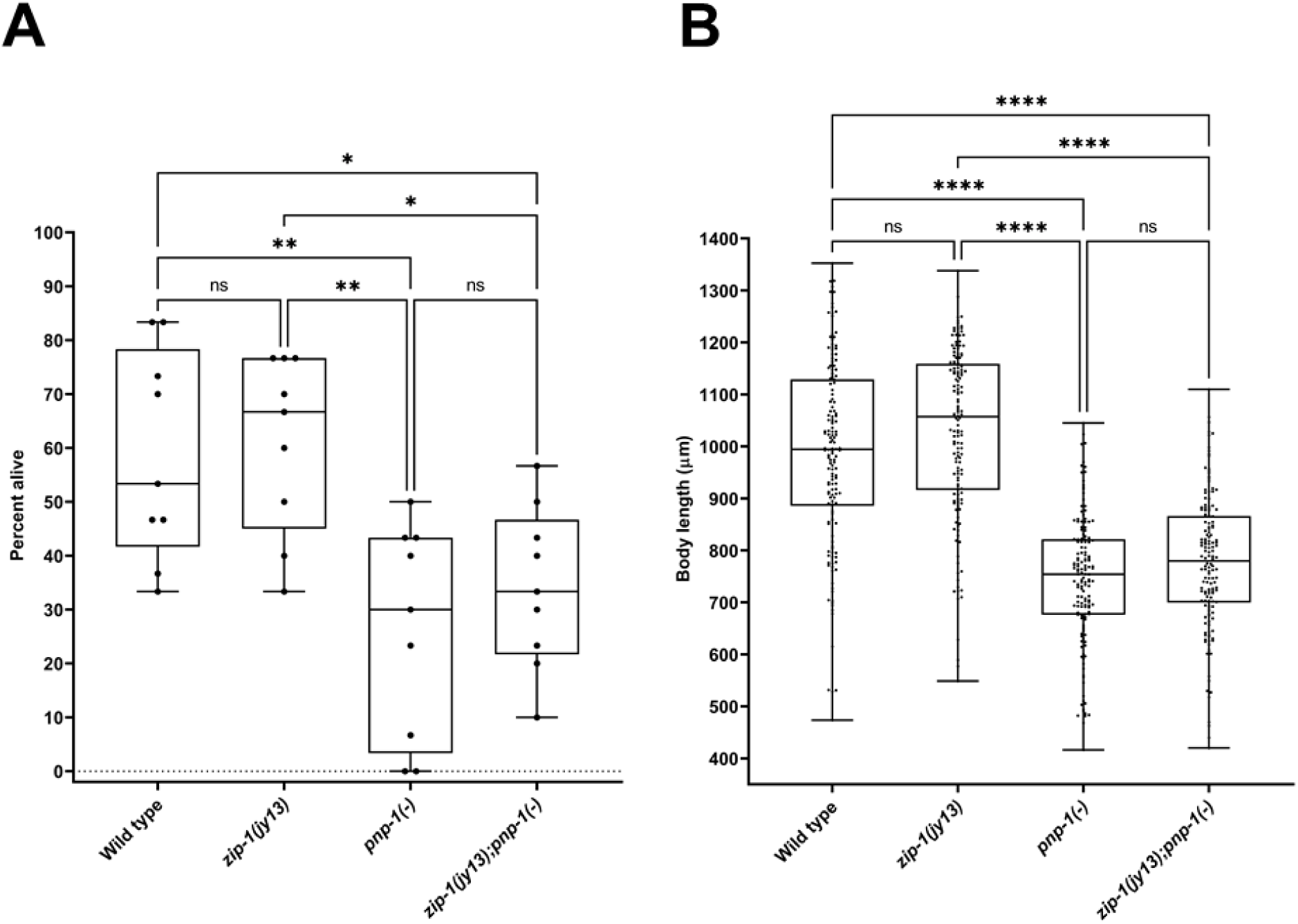
Increased sensitivity to heat shock and smaller size phenotypes of *pnp-1(jy90)* mutants do not depend on *zip-1*. (A) Graphical representation of survival after heat shock. Nine biological replicates from three experiments are indicated with circles; 30 animals were analyzed in each replicate. (B) Graphical representation of body length measurements after 44 h incubation at 20°C. 150 animals were analyzed, 50 animals per each of three replicates. (A, B) The box-and-whisker plots were used for data representation. Each box represents 50% of the data closest to the median value (line in the box). Whiskers span the values outside of the box. All strains are in *jyIs8[pals-5p::gfp; myo-2p::mCherry]* strain background. A Kruskal-Wallis test was used to calculate *p*-values; **** *p* < 0.0001; ** *p* < 0.01; * 0.01 < *p* < 0.05; ns indicates nonsignificant difference (p > 0.05).

